# Extended Shine-Dalgarno motifs govern translation initiation in *Staphylococcus aureus*

**DOI:** 10.1101/2025.06.19.660503

**Authors:** Maximilian P. Kohl, Roberto Bahena-Ceron, Béatrice Chane-Woon-Ming, Maria Kompatscher, Matthias Erlacher, Pascale Romby, Bruno Klaholz, Stefano Marzi

## Abstract

Regulation of translation initiation is central to bacterial adaptation, but species-specific mechanisms remain poorly understood. We present high-resolution mapping of translation start sites in *S. aureus,* revealing distinct features of initiation alongside numerous unannotated small ORFs. Our analysis, combined with cryo-EM of a native mRNA-ribosome complex, shows that *S. aureus* relies on extended, start codon proximal Shine-Dalgarno (SD) interactions, creating specificity against phylogenetically distant bacteria. Several natural *S. aureus* initiation sites are not correctly decoded by *E. coli* ribosomes. We identify new and conserved non-canonical start codons, whose regulatory initiation sites contain these characteristic extended SD sequence motifs. Finally, we characterize a striking example of uORF-mediated translational control in *S. aureus*, demonstrating that translation of a novel small leader peptide modulates expression of a key biofilm regulator. The described mechanism involves codon rarity, ribosome pausing and arginine availability, linking nutrient sensing to biofilm formation in this major human pathogen.

Graphical Abstract

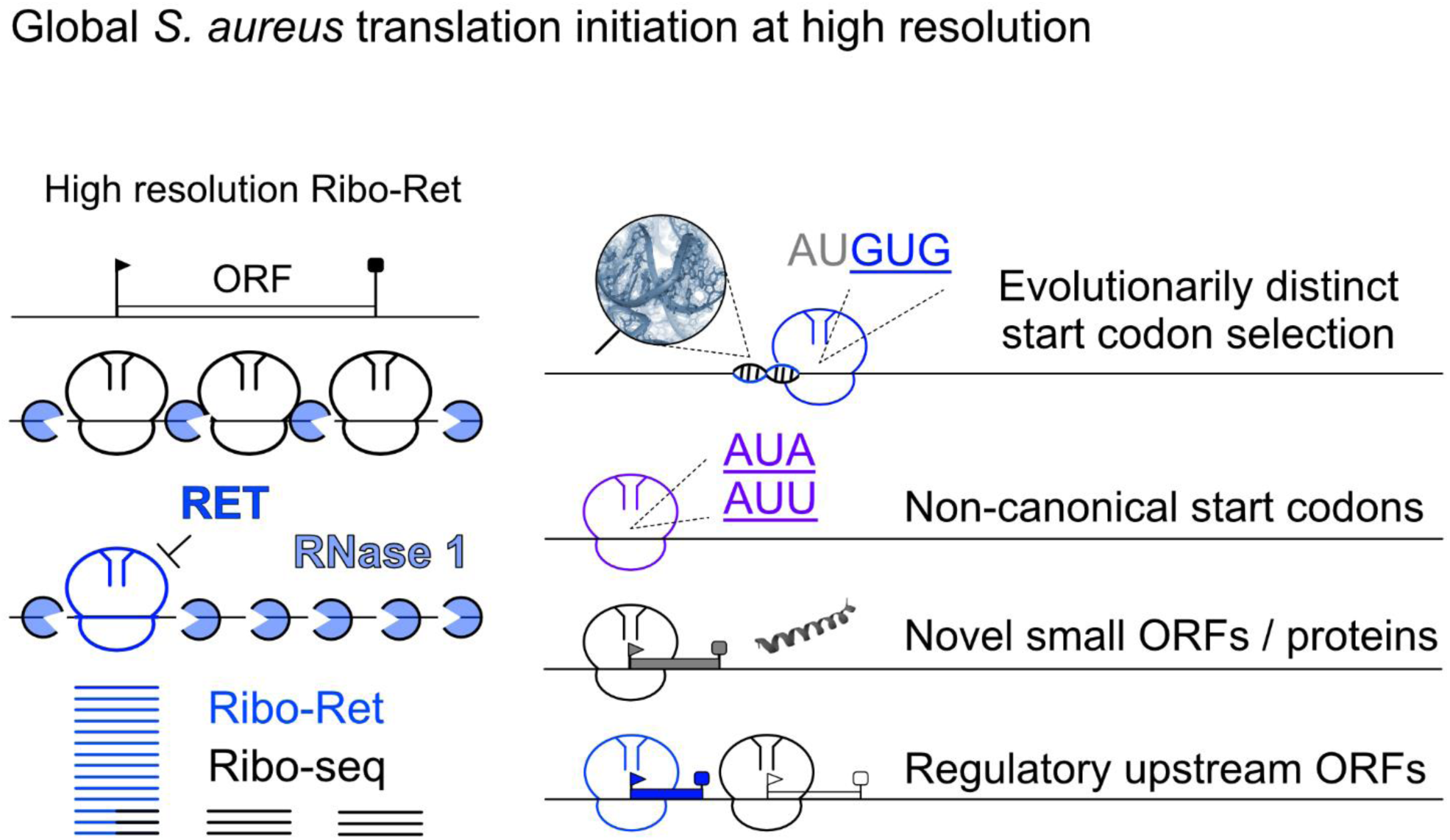

## Introduction

*Staphylococcus aureus* is a major opportunistic pathogen responsible for a wide range of community and hospital acquired infections. To survive in the host and ensure successful dissemination, it has evolved complex regulatory networks that respond to environmental cues by modulating the expression of numerous virulence factors (1). While transcriptional regulation and sRNA-mediated control have been well characterized (2, 3), alternative mechanisms acting at the level of translation remain largely unexplored. In bacteria, one of the key targets for regulation is the rate-limiting translation initiation step (4), which can be rapidly modulated in response to environmental cues and stress. Canonical translation initiation begins with the binding of the small ribosomal subunit (30S) to a translation initiation site (TIS) on a given mRNA. Within the TIS, the correct open reading frame (ORF) is defined by the presence of a start codon, usually AUG, GUG or UUG, which is decoded by formyl-methionyl initiator tRNA^fMet^ in the ribosomal P-site. Start codon recognition represents a key step, resulting in a conformational change of the 30S initiation complex, enabling joining of the large ribosomal subunit and transitioning into the elongation phase of protein synthesis (5, 6). While both the efficiency as well as the accuracy of this decoding step are modulated by the presence of initiation factors (5–7), the initial ribosome recruitment is predominantly governed by the biophysical properties of an mRNA (8, 9). The accessibility of the TIS, particularly the presence of unstructured regions, plays a central role (9–11). To facilitate and enhance ribosome binding, many mRNAs utilize Shine-Dalgarno (SD) sequence motifs, complementary to the 3′ end of the 16S rRNA, designated anti-SD (aSD) (12). The SD-aSD interaction forms a short RNA helix at the ribosomal platform (13, 14), where several ribosomal proteins may further modulate mRNA binding (4). For example, *Escherichia coli* ribosomal protein bS1 is known to mediate affinity towards single-stranded mRNA regions (15, 16) and contributes to the unwinding of secondary structures near the TIS (10). Extensively studied in *E. coli*, the SD-aSD interaction is not strictly required for correct start site selection (17–19) but is thought to play a supportive role (17, 20), although it can influence start codon recognition once the TIS is determined (21, 22). While our current understanding of bacterial translation initiation is largely based on studies in *E. coli*, growing evidence points to a functional diversification across bacterial species. Differences in SD usage, transcriptome architecture, rRNA sequence and ribosomal protein composition suggest species-specific tuning of translation initiation mechanisms (18, 23, 24). Of particular note, differences in the nature of ribosomal binding sites (RBS) have been recognized (25–27), but the underlying molecular basis remains unknown.

Precise mapping of translation initiation sites is vital to decipher the mechanistic divergence across bacteria, and to uncover the full coding potential of their genomes. However, small ORFs (sORFs) can elude conventional annotations and detection by proteomics (28–30). Ribosome profiling (Ribo-seq) is the state-of-the-art technique to monitor translation dynamics globally (31), but its resolution in bacteria has historically lagged behind that achieved in eukaryotes. In eukaryotic systems, the use of RNase 1 for the generation of ribosome protected fragments (RPFs), in combination with harringtonine, enabled high-resolution mapping of initiation sites (32). This approach has led to key discoveries such as the widespread translation of upstream ORFs (uORFs) functioning as regulators, which respond to stress and signaling pathways (32, 33). Furthermore, it led to the identification of several start sites utilizing non-canonical initiation triplets (34, 35) and provided valuable insights into alternative initiation mechanisms and isoform diversity (36). In bacteria, the main limitations stem in part from technical constraints in RPF generation. In comparison, RNase 1 has been underused due to its inactivity in *E. coli* extracts (37) and is typically replaced by micrococcal nuclease (MNase) (20, 38–44), whose strong sequence bias limits obtainable resolution (45, 46). However, a recent study has shown that RNase 1 can generate high-resolution RPFs in *Listeria* and *Salmonella* (47). Therefore, we employed RNase 1 in *Staphylococcus aureus*, significantly enhancing reading frame resolution. We further coupled its use with the initiation-specific inhibitor Retapamulin (Ribo-Ret), which has been used for global mapping of initiation sites in *E. coli* (48).

Our study now analyzes precise start codon selection on a translatome-wide scale in *S. aureus*, revealing accurate decoding of multiple sites, which could not be rationalized by the current understanding of bacterial translation initiation. Functional analysis shows that *E. coli* ribosomes fail to recognize the correct start codon when presented with specific natural *S. aureus* mRNAs, underscoring mechanistic differences in start site selection. A structural analysis by cryo electron microscopy (cryo-EM) of the *S. aureus* ribosome in complex with a native *S. aureus* mRNA reveals start-codon proximal extended SD-aSD interactions. Such rigid SD-aSD helix formation on the platform of the 30S ribosomal subunit limits the flexibility observed in *E. coli* ribosomes, directly influencing start codon recognition. This new mechanistic understanding led us to assess novel candidate initiation sites in *S. aureus* with greater confidence and uncovered a diverse set of previously unrecognized regulators. Among these is the use of broadly conserved non-canonical start codons, which reduce initiation efficiency. The corresponding mRNAs all feature extended SD sequences, which aid recognition of the weak start codons. We also present the first comprehensive mapping of sORFs, including putative regulatory uORFs in *S. aureus*, and uncover a striking example of conditional regulation mediated by an unannotated leader peptide. Translation of this peptide directly influences the expression of the downstream transcription factor Rbf, a key regulator of biofilm formation (49). Its uORF-dependent regulation is driven by a combination of codon rarity and arginine availability, leading to ribosome pausing, consequent occlusion of the *rbf* ribosome binding site, and inhibition of translation initiation under arginine-limiting conditions. Collectively, our findings offer a precise mechanistic understanding of the pronounced differences in translation initiation, driven by the evolution of alternative RBS architectures across bacteria, and offer a broad perspective on translational control in a clinically relevant pathogen.

## Results

### RNase 1 Ribo-Ret reveals global *S. aureus* translation initiation at high resolution

To capture a comprehensive snapshot of the *S. aureus* translatome, cells were grown towards the mid-to late exponential growth phase in rich medium, where virulence factors are highly expressed. Brief treatment of the cultures with Retapamulin (Ret) resulted in a pronounced arrest of initiating ribosomes (Figure 1A, B). This was carefully monitored with non-digested controls, whose polysome profiles were analyzed in parallel (Figure 1B). The addition of Ret resulted in a stark shift in the sucrose gradient from the polysome-towards the 70S fraction, indicating stalling during the initiation step. Likewise, treatment of the cell extracts with either MNase or RNase 1 enriched 70S peaks and resulted in a loss of polysomes, demonstrating a successful digestion for the generation of RPFs (Figure 1B). Both MNase and RNase 1 digested samples, Ret-treated and non-treated, were processed for RPF-library preparation and next-generation sequencing.

**Figure 1.**
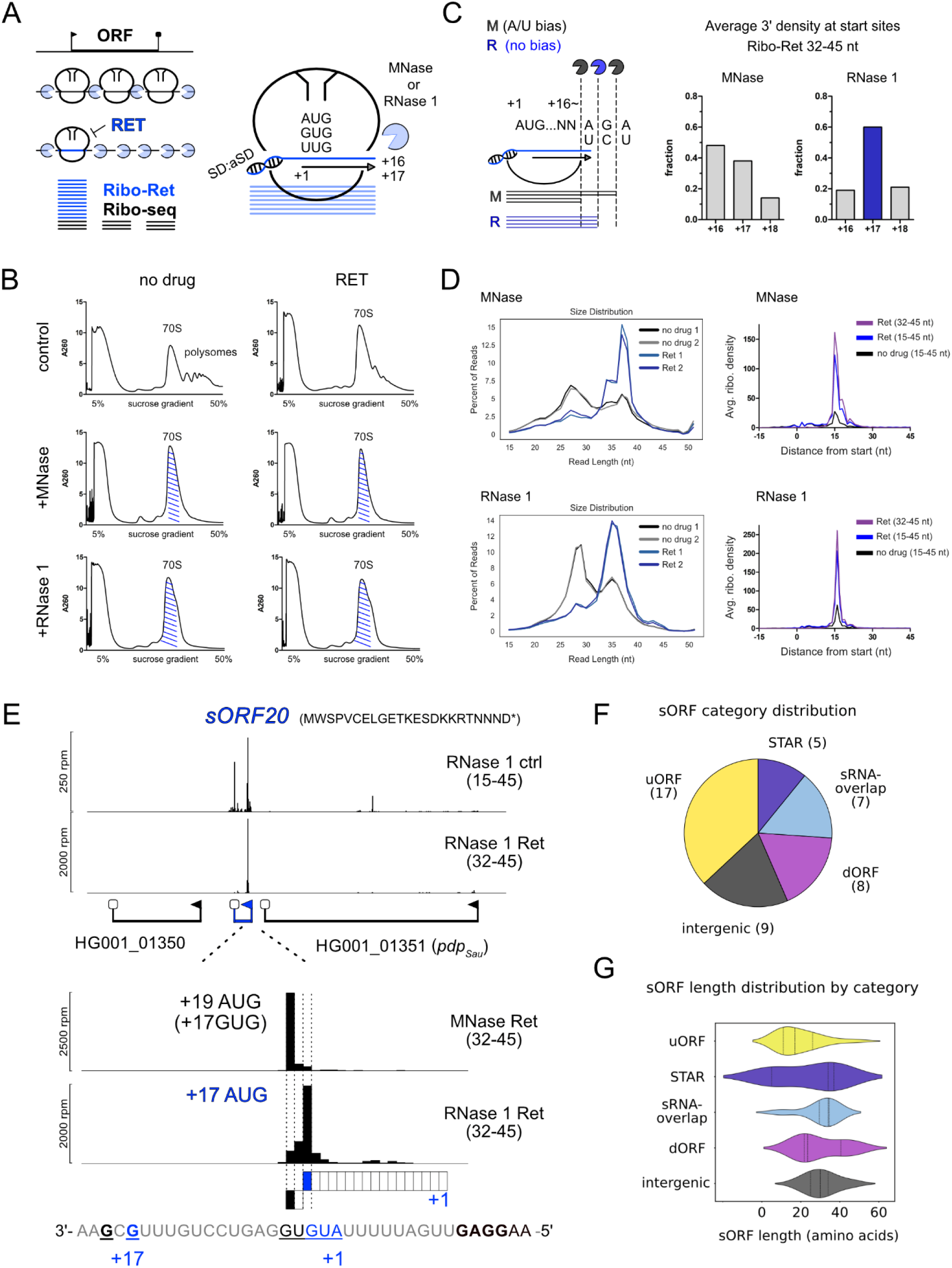
RNase 1 Ribo-Ret enables high-resolution translation initiation profiling and sORF discovery in *S. aureus*. (A) Schematic representation of Retapamulin-assisted ribosome profiling with RNase 1. (B) Sucrose density gradient profiles for Ribo-Ret with differential nuclease treatment and mock treated controls. Monosome fractions isolated and processed are highlighted in blue. (C) Schematic depiction of MNase cleavage bias and the obtained improved resolution by RNase 1 Ribo-Ret. Fractions indicate the relative amount of mapped read 3’ ends distributed between positions +16, +17 and +18 in the peak window of a global average gene analysis. (D) Read length distribution of Ribo-seq (no drug) and Ribo-Ret (Ret) experiments obtained with either nuclease as well as plots of the average gene analysis used to assess start site resolution. Ret treated samples are highlighted in blue in both panels and in purple for the selective analysis of longer reads on the average gene plots. (E) Exemplary depiction of sORF discovery and start codon assignment in form of Ribo-seq and Ribo-Ret profiles of *sORF20*. Sample type and considered read lengths are indicated accordingly. For example, an RNase 1 untreated ctrl Ribo-seq profile is shown, considering reads of 15-45 nt in length. The annotation designates RNase 1 ctrl (15–45). (F, G) Newly identified sORFs by category and their comparative sORF length distributions by kernel density violin plots. Also see Figure S1, S2 and Table S1.

Sequencing results were first analyzed to assess Ribo-seq quality to see whether RNase 1 could improve the resolution of Ribo-Ret in *S. aureus* (Figure 1C, D). For either nuclease, in the absence of Ret, the read lengths formed a bimodal distribution with peaks at approximately 26-29 nt and 35-38 nt, respectively. Strikingly, the addition of Ret resulted in a strong shift towards the longer read population (Figure 1D). The observed distribution is consistent with the approximately 28 nt protection from nuclease digestion, conferred by elongating ribosomes. The stark increase in read length upon Ret treatment can be rationalized by the additional protection that SD-aSD base pairings confer during initiation (50).

We next performed an average gene analysis of 3’ end densities, mapped across annotated open reading frames, and confirmed a drastic enrichment of densities corresponding to initiating ribosomes in the Ret-treated samples (Figure 1D). The peak positions, relative to annotated start codons, mapped at the expected 16 nt distance from the first nucleotide of the start codon (+17). As hypothesized, these peaks were sharper for RNase 1 and mapped predominantly (60%) to this position (Figure 1C). This was in contrast to MNase, where preferential cleavage bias resulted in a broader distribution. Hence, RNase 1 Ribo-Ret provides global and high-resolution translation initiation snapshots.

### Discovery of new sORFs by high-resolution Ribo-Ret

While standard Ribo-seq data can be used to evaluate potential sORFs, the application of Ribo-Ret and the observed enrichment in ribosome density at putative start sites, provides stronger experimental support. We therefore leveraged our complementary Ribo-seq and Ribo-Ret datasets for comprehensive small ORF discovery (Figures 1E, F, G as well as S1 and S2). The pipeline used for the discovery of the sORFs is described in the Supplementary Material (Scheme S1). All considered ribosome densities were consistent between the two complementary datasets and RNase 1 allowed precise start codon assignment.

The obtained improved resolution was critical when Ribo-Ret densities could match to multiple potential sORFs. This was particularly the case for overlapping start codon sequences (e.g., AUGUG). A typical example is shown in Figure 1E, where Ribo-Ret density mapped to a newly identified sORF (*sORF20*). In the RNase 1 dataset, the 3’ ends of Ribo-Ret reads accurately mapped at position +17 to an AUG start codon, while in the MNase dataset the corresponding peak falsely indicated translation from an overlapping GUG start codon. In-frame AUG start codon selection on *sORF20* mRNA was further biochemically validated by toeprinting of an *S. aureus* 70S initiation complex stalled with Ret (Figure S1).

Focusing our analysis on intergenic regions, we obtained a high confidence sORF map specific to our experimental growth conditions (Table S1). In addition to previously identified functional small proteins (e.g., phenol soluble modulins (51)), our study led to the discovery of 46 unannotated sORFs. To our knowledge, only two of these had previously been identified in a proteomics study (52), and one additional sORF had been reported to function in transcription antitermination within the isoleucine biosynthesis operon (53). The other 43 sORFs represent *bona fide* novel candidates. Notably, several of these sORFs are predicted to encode small hydrophobic transmembrane helices, including *sORF26* and *sORF36*. Through toeprinting assays, analyzing *S. aureus* 30S initiation complex formation, their efficient ribosome recruitment was biochemically validated (Figure S1C, D). Overall, 14 initiation sites were analyzed by toeprinting, confirming active start site selection (Table S1).

Based on the genomic context, identified sORFs were classified into distinct categories (Figure 1F). Several examples overlapped with previously annotated putative sRNAs features (s*ORF6, 26, 30, 34, 35, 36 and 40)* and were therefore assigned an ‘sRNA overlap’ category. We further identified nine sORFs located in intergenic regions, which were distinguished from sORFs embedded within 5’-or 3’ “untranslated” regions, designated as upstream ORFs (uORFs) or downstream ORFs (dORFs), respectively. Among them, 17 sORFs could be classified as uORFs and 8 as dORFs. The *sORF20* (Figure 1E), while possessing its own promoter, was categorized as dORF because it can be co-transcribed with its immediate upstream gene *Pdp_Sau_* (HG001_01351), which encodes a recently identified anti-phage defense protein (54). To complete our sORF categorization, we observed that several of the newly identified initiation sites shared sequence similarity and conserved ribosomal binding sites (RBSs). These motifs stem from the conserved 5’ region of *Staphylococcus aureus* repeat (STAR) motifs (Figure S2). We conclude that STAR motifs consistently recruit ribosomes when expressed. Among the 46 sORFs identified in our study, intergenic sORFs, dORFs and sRNA overlapping sORFs tend to be longer than uORFs, which may act as regulatory leader peptides (Figure 1G). In contrast, STAR-associated sORFs show considerable length variability (Figure 1G).

### Translation initiation in *S. aureus* follows distinct rules to define start codon selection

Our high-resolution RNase 1 Ribo-Ret analysis facilitated correction and re-annotation of approximately 40 coding sequences. These corrections ranged from minor adjustments, with accurate assignment of start codons among multiple in-frame possibilities, to more substantial changes as illustrated for *sarX* (Figure S3). Importantly, while manually inspecting translation initiation sites, we observed unexpected patterns in start codon selection. Specifically, our RNase 1 data revealed cases of precise start codon definition despite potential ambiguity and competition. A typical example is the aureolysin (*aur*) mRNA, whose GUG start codon overlaps with an alternative AUG start codon in an AUGUG context, yet its correct initiation triplet is consistently selected (Figure 2A, B). This was unexpected, as studies in *E. coli* have shown that a relatively shorter spacer length between the SD and the start codon is generally preferred (21), making the AUG in this context seemingly better positioned for initiation. Using our improved HG001 annotation, we analyzed spacing preferences and observed that on average, the aligned spacing between SD and start codon in *S. aureus* is increased by two to three nucleotides compared to *E. coli* (Figure 2A). Although variations in spacer lengths among bacteria have previously been observed (25), they have not been linked to differences in start codon selection.

**Figure 2.**
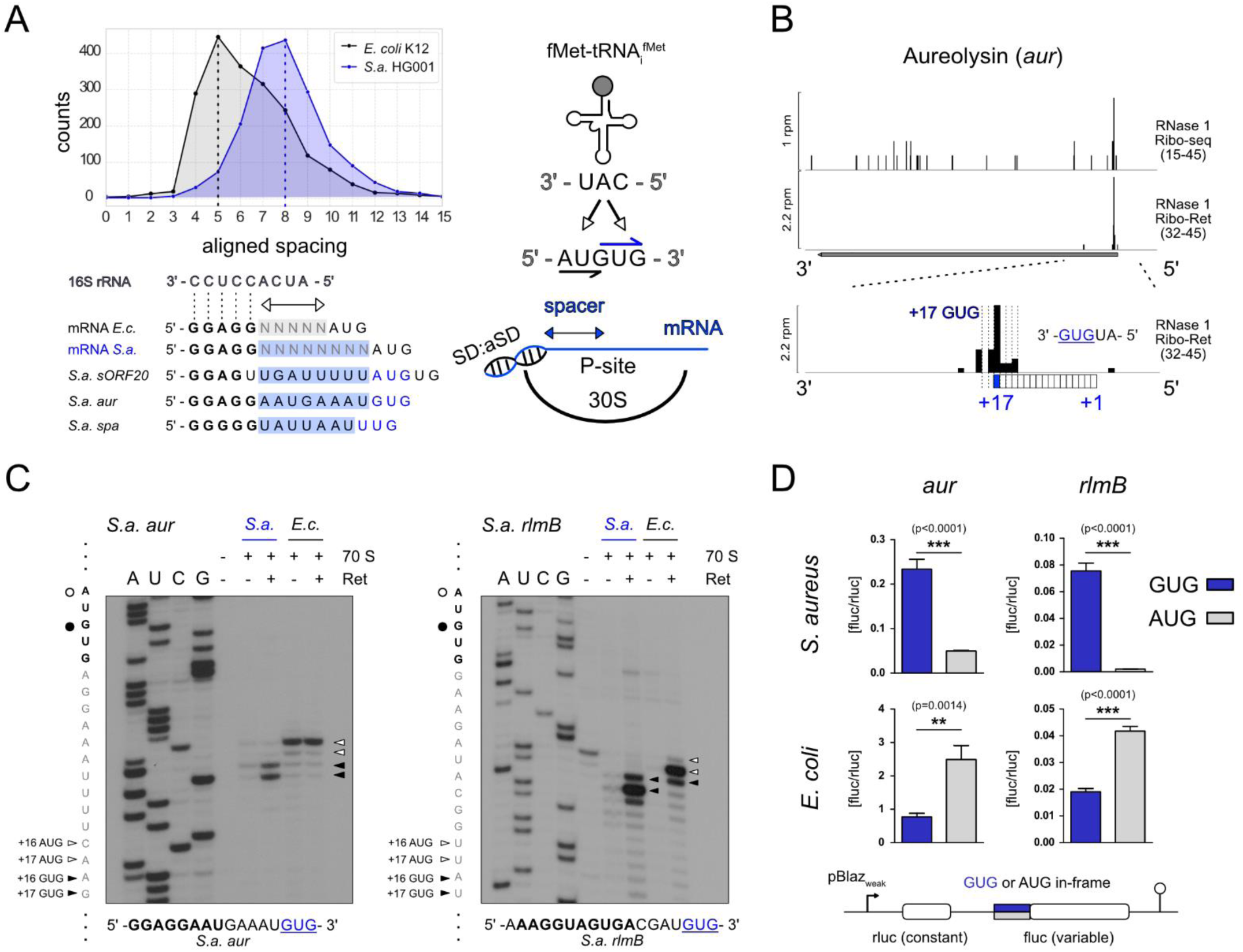
High resolution Ribo-Ret reveals evolutionarily distinct start codon selection. (A) Comparative analysis of aligned spacing distributions between *E. coli* and *S. aureus*. Exemplary *S. aureus* ribosome binding sites are displayed for reference and their in-frame start codons are highlighted in blue along a schematic representation of SD-aSD directed localization of the start codon in the P-site. Also see Figure S3. (B) Ribo-seq and Ribo-Ret profiles of *S. aureus aur* mRNA showing ribosomal density peaks, highlighting selection of an in-frame GUG start codon within an AUGUG overlap. (C) Toeprinting analysis of natural *S. aureus* mRNAs *aur* and *rlmB* decoded by *S. aureus* (*S. a*) and *E. coli* (*E. c*) 70S ribosomes. Sequencing lanes (A, U, C, G) and toeprint positions (arrows) are indicated accordingly. Ret denotes Retapamulin. (D) Results from *in vivo* dual luciferase assays measuring translation initiation efficiencies from in-frame AUG or GUG reporter fusions of *aur* and *rlmB* initiation sites in *S. aureus* and *E. coli*. Measurements corresponding to correct decoding of the natural in-frame GUG start codon are highlighted in blue. A schematic drawing of the dual luciferase reporter is depicted. Mean and standard deviation from at least five biological replicates are shown. Two-tailed unpaired t test, ∗∗∗p < 0.001.

Because the identified initiation sites, such as *aur*, imply differences in the bacterial translation initiation mechanism, we hypothesized that several natural *S. aureus* mRNAs may not be accurately decoded by *E. coli* ribosomes. To test this hypothesis, we performed cross-species experiments and studied start codon selection in the context of the *S. aureus aur* and *rlmB* initiation sites, which feature AUGUG overlaps and require recognition of an in-frame GUG start codon. Using purified *E. coli* or *S. aureus* 70S ribosomes supplied to a recombinant *in vitro* translation system (see Methods), we analyzed start codon usage by toeprinting of initiation complexes stalled in the presence of Ret. Consistent with our hypothesis, *E. coli* 70S preferentially selected the AUG positioned proximal to the SD helix, while only *S. aureus* ribosomes accurately decoded the in-frame GUG start codon (Figure 2C), suggesting species-specific initiation preferences. To further validate these results *in vivo*, we constructed bi-cistronic dual reporter plasmids, in which either the AUG or GUG start codon of *aur* and *rlmB* was fused in-frame to a Firefly luciferase gene. The expression was normalized to the co-transcribed Renilla luciferase with a constant translation initiation context. Our data demonstrate that *S. aureus* strongly favors the annotated GUG start codon, whereas *E. coli* shows significantly higher expression from AUG *in vivo* (Figure 2D).

### Extended SD-aSD base pairing confers evolutionarily distinct start codon selection

To further investigate differential start codon selection, we performed toeprinting on *spa* mRNA, a key *S. aureus* transcript that encodes the virulence factor protein A. Specifically, we altered the native AU<u>UUG</u> start codon context of *spa* mRNA by a single nucleotide change to create an AU<u>GUG</u> overlap. Consistent with the results of *aur* and *rlmB* mRNAs, this mutant *spa* mRNA was decoded differentially by *S. aureus* and *E. coli* (Figure 3A). *S. aureus* ribosomes decoded the in-frame GUG start codon while *E. coli* ribosomes ambiguously selected both the AUG and the GUG start codons in this sequence context.

**Figure 3.**
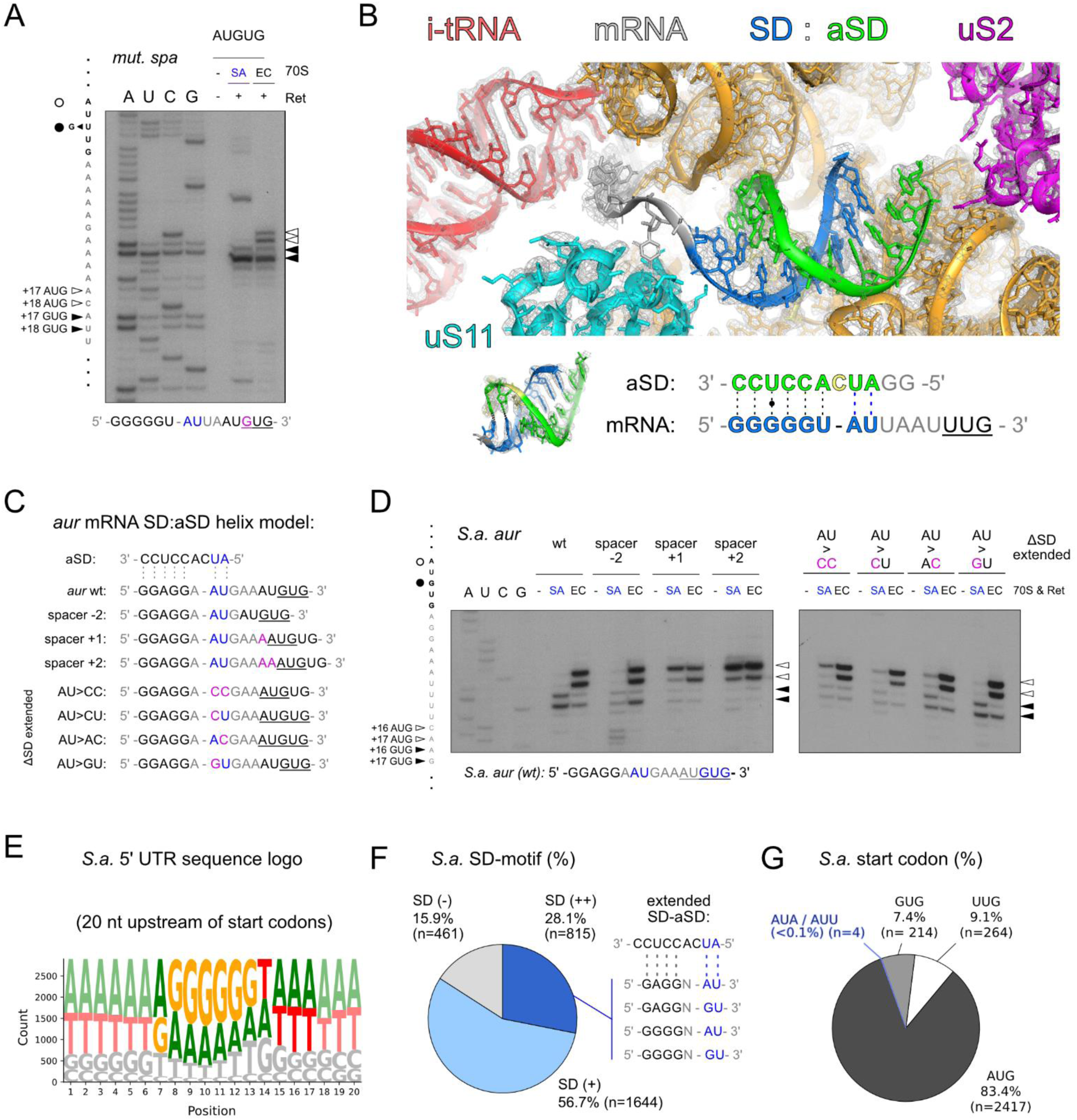
Cryo-EM structure unveils an extended SD-aSD helix directing start codon selection. (A) Toeprinting analysis of mutant *spa* mRNA with ambiguous translation initiation context, decoded by *S. aureus* or *E. coli* 70S ribosomes. Sequencing lanes (A, U, C, G), toeprint positions (black and empty arrows), and the exact RBS sequence are shown. (B) Cryo-EM map and model of the extended SD-aSD helix region within an *S. aureus* 70S initiation complex formed on the natural *spa* mRNA. The corresponding sequence is shown, highlighting nucleotides involved in Watson-Crick base pairing. Initiator tRNA is shown in red, the aSD sequence is displayed in green, the *spa* mRNA’s SD motif is colored in blue and the mRNA’s model towards the start codon shown in gray. Start codon proximal extended interactions are shown from an alternative angle on the bottom left. (C, D) Comparative *S. aureus* and *E coli* 70S toeprinting analysis of *aur* mRNA sequence variants, analyzing the contribution of spacer length and extended SD-aSD base pairing to the decoding preferences of its natural AUGUG start codon overlap. Predicted extended SD-aSD pairing interactions are shown in C, while the toeprinting results are displayed in panel D. mRNA sequencing lanes (A, U, C, G) are indicated and the respective toeprint positions highlighted by black and empty arrows. Also see Figure S4. (E) Global sequence motif analysis of *S. aureus* RBS sequences upstream of initiation sites. (F) Computationally predicted SD sequence usage among *S. aureus* mRNAs and the percentage utilizing extended SD-aSD base pairing interactions. (G) Global start codon usage percentage among annotated *S. aureus* mRNAs.

To better understand the mechanistic details of *S. aureus* translation initiation, we determined the cryo-EM structure of an *S. aureus* 70S ribosome initiation complex programmed with native *S. aureus spa* mRNA. The structure shows a characteristic SD-aSD helix and allows to closely examine the interactions, which are critical for accurate start codon positioning in the ribosomal P-site. Strikingly, beyond the canonical SD-aSD helix typically seen in bacteria (13, 14), two additional Watson-Crick base pairs are found (Figure 3B). These base pairs form between an AU dinucleotide following the canonical SD and nucleotides A1543 and U1544 on the 3’-region of the 16S rRNA. This extended duplex occupies a broader region of the ribosomal platform, spanning further along the mRNA path, from uS11 toward uS2 (Figure 3B). The additional AU dinucleotide corresponds to the increased spacing between SD and start codon in *S. aureus* that we found (Figure 2A), and strengthens the SD-aSD interactions, i.e. the sequence insertion is not a linker increase but rather it provides additional complementary base pairing interactions.

To assess whether this extended SD-aSD base pairing could resolve start site ambiguity in natural sequence contexts, we performed toeprinting on variants of the previously analyzed *aur* mRNA. Based on the extended Watson-Crick base pairing seen in the *spa* RBS, we predicted similar interactions for the *aur* RBS (Figure 3C). We first confirmed that *aur* start codon selection by *S. aureus* 70S was dependent on spacer length. Indeed, a single nucleotide insertion shifted decoding from the in-frame GUG to the competing AUG start codon, without affecting the *E. coli* ribosome’s recognition of AUG (Figure 3D). We next introduced mutations altering the predicted start codon-proximal AU dinucleotide involved in the extended SD-aSD pairing (Figure 3A, D) to study its contribution to start codon definition. A substitution with a CC pair abolished the GUG start codon recognition, and a CU dinucleotide failed to restore it. Partial recovery was observed with AC, while full restoration occurred with a GU pair, indicating that a G:U wobble pair can substitute for A:U Watson-Crick interactions to maintain accurate start codon selection (Figure 3D). *E. coli* 70S remained unaffected by any of these changes and consistently decoded the AUG initiation triplet.

The observed accessibility of nucleotides upstream of the canonical CCUCC sequence in the 3′ region of *S. aureus* 16S rRNA enables new possibilities for SD-aSD interactions and may additionally contribute to accurate start site selection on natural mRNAs. A typical example is the *S. aureus rarA* mRNA, which contains a potentially ambiguous AUGUG start site but is accurately decoded despite a poorly defined canonical SD sequence (Figure S4). We hypothesized that, in such contexts, alternative energetically favorable SD-aSD duplexes could guide correct start codon placement in the P-site. This is supported by 30S toeprinting, which showed accurate GUG selection by *S. aureus* ribosomes, while *E. coli* 30S failed to efficiently decode either start codon (Figure S4A). Because all studied examples are in the context of AUGUG overlaps, we analyzed one further example with two competing AUG start codons, designating alternative reading frames. Such an arrangement is found, for example, in the natural *proP* mRNA of *S. aureus,* where precise start codon selection is observed by Ribo-Ret and predicted to depend on extended SD-aSD pairing (Figure S4B). Consistent with our hypothesis, *S. aureus* 30S accurately decoded the second AUG start codon, whereas initiation complex formation was ambiguous between the two start sites for *E. coli*. Jointly, these findings indicate the important contribution of extended SD-aSD formation to start codon definition during translation initiation in *S. aureus*.

Interestingly, most *S. aureus* mRNAs feature AU-rich spacer sequences, which may similarly promote extended SD-aSD interactions, as revealed by global sequence motif analysis (Figure 3E). While we predict the presence of canonical SD-sequences in approximately 84% of *S. aureus* mRNAs based on computational SD-motif search, our analysis indicates a conservative estimate of approximately 30% of initiation sites relying on extended SD-aSD base pairing (Figure 3F). This prediction is based on the experimentally validated base pairing possibilities of the natural *aur* and *spa* mRNAs and does not include other alternative interactions, which cannot be excluded at this stage. Finalizing our computational analysis of *S. aureus* translation initiation contexts, we report an unusually high frequency of UUG start codons, approximately 9.1% of all start sites, compared to around 2% in *E. coli* (Figure 3G). In *S. aureus*, the remaining start codons correspond to 83.4% to AUG and to 7.4% to GUG. Strikingly, four initiation sites designated non-canonical start codons, which we further analyzed (Figure 4 and S5).

**Figure 4.**
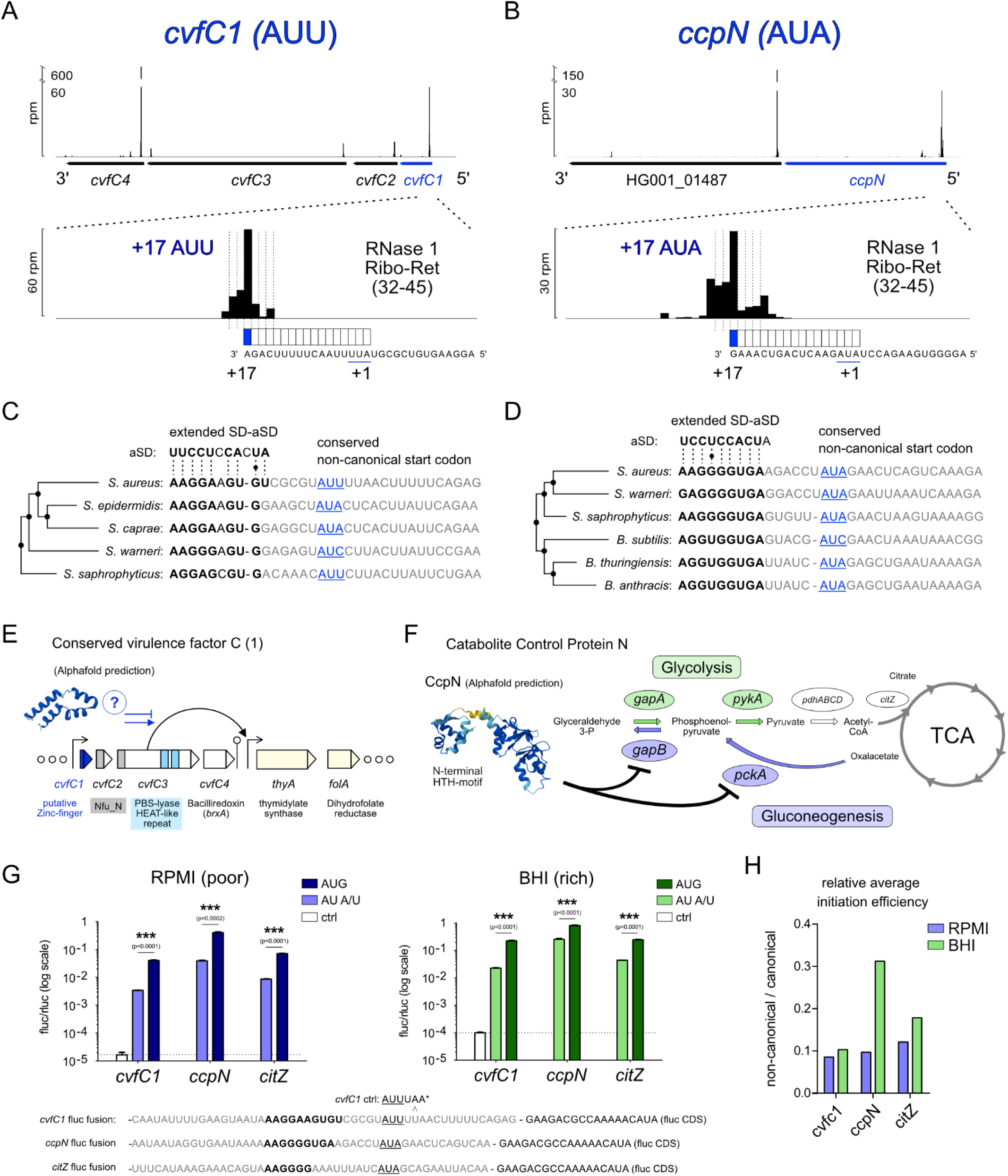
Ribo-Ret facilitates discovery of conserved non-canonical start codons. (A, B) Ribo-Ret profiles of *S. aureus cvfC1* (A) *and ccpN* (B) showing ribosomal density peaks and a zoomed focus on their respective initiation sites, highlighting the use of non-canonical start codons. Also see Figure S5. (C, D) Sequence conservation of *cvfC1* (C) and *ccpN* (D) translation initiation contexts with extended SD-aSD base pairing and non-canonical start codon (in blue characters) selection. (E) Schematic depiction of *cvfC1’s* position in the *cvfC* operon. A putative Zinc finger fold of CvfC1 could indicate a contribution to the transcriptional regulation of *thyA* linked with CvfC3 expression. (F) Schematic depiction of CcpN’s functional role as a transcriptional repressor of gluconeogenesis. CcpN’s predicted structure via AlphaFold is shown and black blunt arrows indicate repression of key gluconeogenic enzymes. (G) Results from *in vivo* dual luciferase assays measuring translation initiation efficiencies from non-canonical (AUU or AUA) and canonical (AUG) start codon reporter fusions of *cvfC1, ccpN* and *citZ* initiation sites in RPMI and BHI media, respectively. Measurements from variable Firefly luciferase reporter fusions are normalized by constant Renilla expression. For *cvfC1,* an additional plasmid variant with in-frame stop codon serves as negative control. Results are displayed on a logarithmic scale as mean and standard deviation from at least three biological replicates. Two-tailed unpaired t test, ∗∗∗p < 0.001. (H) Comparative plot of the average relative initiation efficiencies (fluc/rluc) of measurements displayed in the panel G, comparing non-canonical against canonical start codon usage in RPMI and BHI media.

### Newly identified non-canonical start codons regulate translation initiation in *S. aureus*

Several parameters modulate initiation efficiency including start codon identity. We observed that Ribo-Ret enriched ribosome density at the non-canonical AUA start codon of *infC*, encoding IF3 (Figure S5), which led us to hypothesize that additional examples could be found. Indeed, during HG001 re-annotation, we identified two novel and conserved non-canonical start codons (Figure 4AB). Notably, for all three of these initiation sites, an extended SD sequence motif is predicted to direct the non-canonical start codon selection.

The first new-found site is the non-canonical AUU start codon of *cvfC1,* encoding a 61 amino acid small protein, designated Conserved virulence factor C1. Although little is known about its function, it possesses a potential zinc finger domain and could be involved in transcriptional regulation (Figure 4E). This gene is the first of an operon initially identified in a screen for the discovery of conserved virulence factors (55, 56). We could clearly assign an RNase 1 Ribo-Ret peak to position +17 in respect to the non-canonical AUU start (Figure 4A). Intriguingly, a very strong and extended SD motif (Figure 4C), as well as the non-canonical nature of the start codon are conserved among *Staphylococci*. The second example we identified is the non-canonical AUA start codon of *ccpN*, encoding catabolite control protein N (57). RNase 1 Ribo-Ret density mapped accurately at position +17 (Figure 4B), clearly indicating its expression. Although CcpN’s functional role in gluconeogenesis has only been demonstrated in *B. subtilis*, both a 76% sequence similarity as well as structural similarity according to an AlphaFold prediction, suggest functional conservation (Figure 4F). Consistently, both the non-canonical start codon usage, as well as strong and extended SD-aSD base pairing with appropriate spacer length are conserved between *S. aureus* and *B. subtilis* (Figure 4D).

Our discovery of non-canonical start codon usage suggests that both *cvfC1-* and *ccpN* expression are strongly regulated at the level of translation initiation. To test this hypothesis, we constructed bi-cistronic dual luciferase reporter plasmids, fusing the respective translation initiation context with a firefly luciferase reporter gene and monitored translation efficiencies *in vivo* (Figure 4G). We included the fourth non-canonical start site that we identified, an AUA start codon of citrate synthase *citZ*, conserved in all *S. aureus* strains derived from the NCTC 8325 lineage, commonly used as laboratory strains (Figure S5). Particularly because of CcpN’s role in central metabolism, we analyzed translation initiation in different growth media conditions. As hypothesized, for all three genes, the synthesis of luciferase was strongly decreased by the use of a non-canonical start codon compared to cognate AUG mutants (Figure 4G). For cells grown in poor media (RPMI), all non-canonical start sites conferred approximately 10% of the reporter expression compared to AUG. Surprisingly, we observed that this relative initiation efficiency was increased up to 30% in the case of *ccpN* AUA, when expressed in rich growth medium (BHI; Figure 4G, H). In contrast, we observed only a modest relative increase for *citZ* AUA, and no media dependent effect for *cvfC1* AUU. Therefore, the usage of non-canonical start codons reduced the yield of protein synthesis *in vivo* and particularly sensitized the expression of *ccpN* to metabolic changes.

### Ribo-Ret identified a variety of uORFs representing potential leader peptides

Our high-resolution Ribo-Ret analysis revealed initiation signatures for 17 upstream open reading frames (uORFs) with potential regulatory functions. An overview of these small uORFs and their experimental validation is presented in Figure 5A. To assess their capacity for translation initiation, we performed toeprinting assays on eight selected candidates (Figure 5A-D). Robust 30S initiation complex toeprint signals were observed for seven out of eight tested uORFs, while one candidate yielded only a faint signal. To further validate uORF translation *in vivo*, we generated translational fusions between the full uORF coding sequences and a luciferase reporter gene, constitutively expressed alongside a Renilla luciferase as an internal control. This assay confirmed *in vivo* translation of four uORF candidates. In contrast, the uORF with a faint toeprint signal (*sORF9*) exhibited only low translation efficiency *in vivo* (Figure 5E).

**Figure 5.**
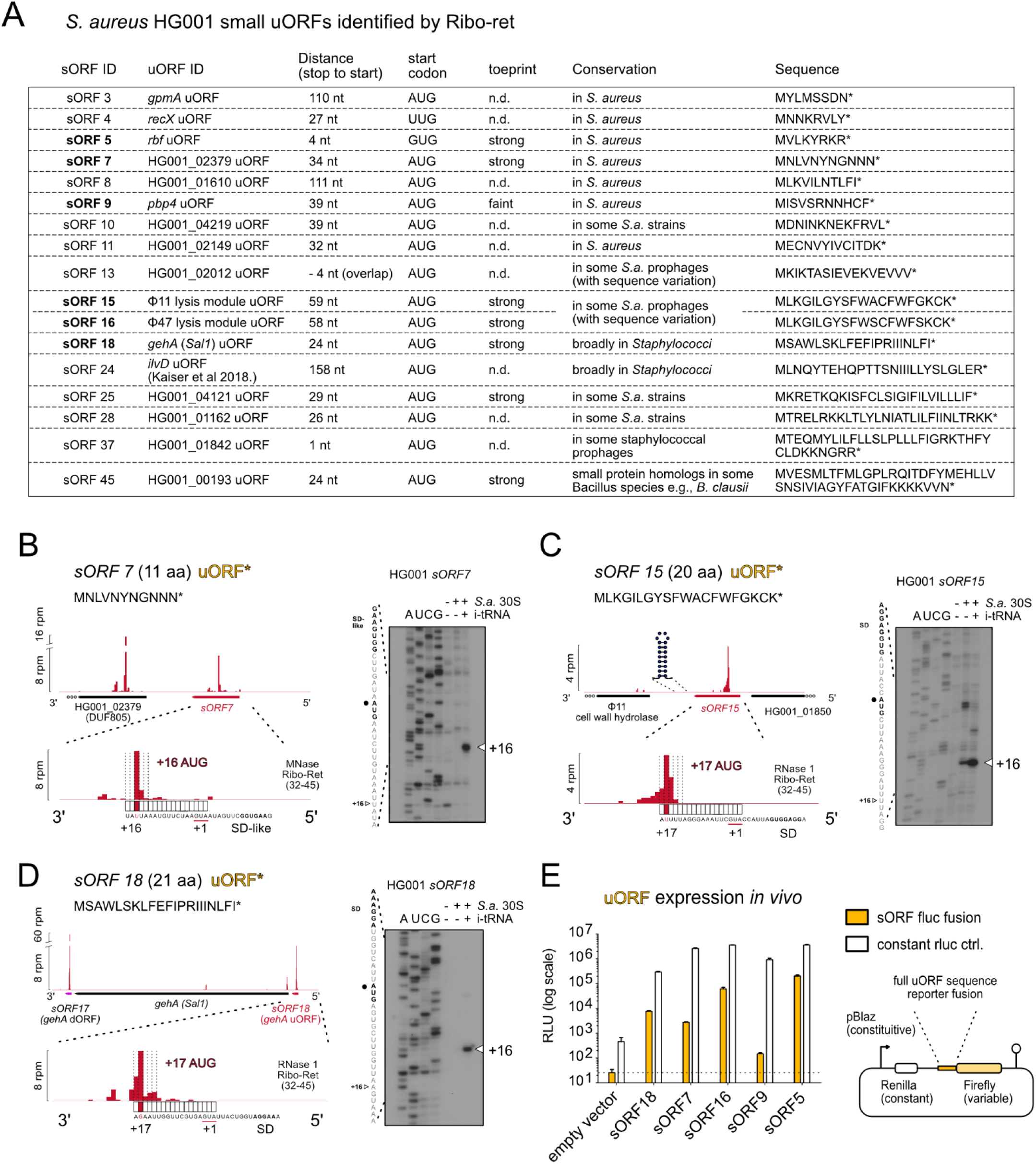
Small uORFs represent candidate regulatory leader peptides. (A) Selected sORF table displaying novel small uORF candidates. Their sORF- and uORF IDs, defined by respective downstream gene annotations, their start codons, coding sequences, distances to downstream genes and conservation are shown. An additional column indicates whether toeprint analyses were performed and if the obtained signal was strong, faint or not determined (n.d.). uORFs that were further experimentally analyzed are highlighted in bold. (B, C, D) Ribo-Ret profiles of *S. aureus sORF7* (B)*, sORF15* (C) and *sORF18* (D), showing ribosomal density peaks and a zoomed focus on their respective initiation sites. Toeprinting analyses validating strong 30S initiation complex formation are shown. The corresponding sequencing lanes (A, U, C, G) and the toeprint positions (+16) are indicated. (E) Selected validation of sORF expression *in vivo* by use of dual luciferase reporter fusions of candidate uORFs, *sORF18, sORF7, sORF16, sORF9 and sORF5*. Firefly luciferase signals from variable sORF reporter fusions are highlighted in orange, whereas measurements from constant Renilla luciferase are shown in white. Absolute measurements (relative light units, RLU) are displayed on a logarithmic scale as mean and standard deviation from at least three biological replicates. A sketch of the bi-cistronic reporter plasmid architecture is shown.

We consider *sORF7* to be an intriguing leader peptide candidate due to its unusually Asn-rich sequence (Figure 5B), but the function of the downstream gene product is unknown. Furthermore, *sORF15* and *sORF16* represent nearly identical uORFs (>90% nucleotide sequence identity), located immediately upstream of the lysis modules within prophages Φ11 and Φ47, respectively (Figure 5C). Notably, both are followed by predicted intrinsic transcription terminators, suggesting a potential cis-regulatory function at the mRNA level. Another interesting discovery is *sORF18,* a 21-amino acid uORF positioned directly upstream of the gene *gehA,* encoding lipase 1. We also note the presence of *sORF17* of similar length, located just downstream of *gehA* (Figure 5D). While this arrangement may reflect independent small protein functions, a potential regulatory role of *sORF18* in *gehA* expression cannot be excluded.

### A small uORF acts as a leader peptide and senses arginine restriction to control *rbf* translation

Short uORFs represent intriguing candidate leader peptides. We hypothesized that in some cases, their translation could act as a sensor for environmental cues or stresses, thereby regulating downstream gene expression. In addition to the Asn-rich *sORF7* (Figure 5B), one uORF in particular (*sORF5*) drew our attention due to its location immediately upstream of *rbf* (**r**egulator of **b**iofilm **f**ormation) (Figure 6A). Notably, the luciferase reporter fusion of *sORF5* showed the highest *in vivo* expression levels among all tested candidates (Figure 5E). Given its close proximity to and partial overlap with the *rbf* RBS, we investigated its potential function as regulatory leader peptide, hereafter referred to as *rbfL* (Figure 6).

**Figure 6.**
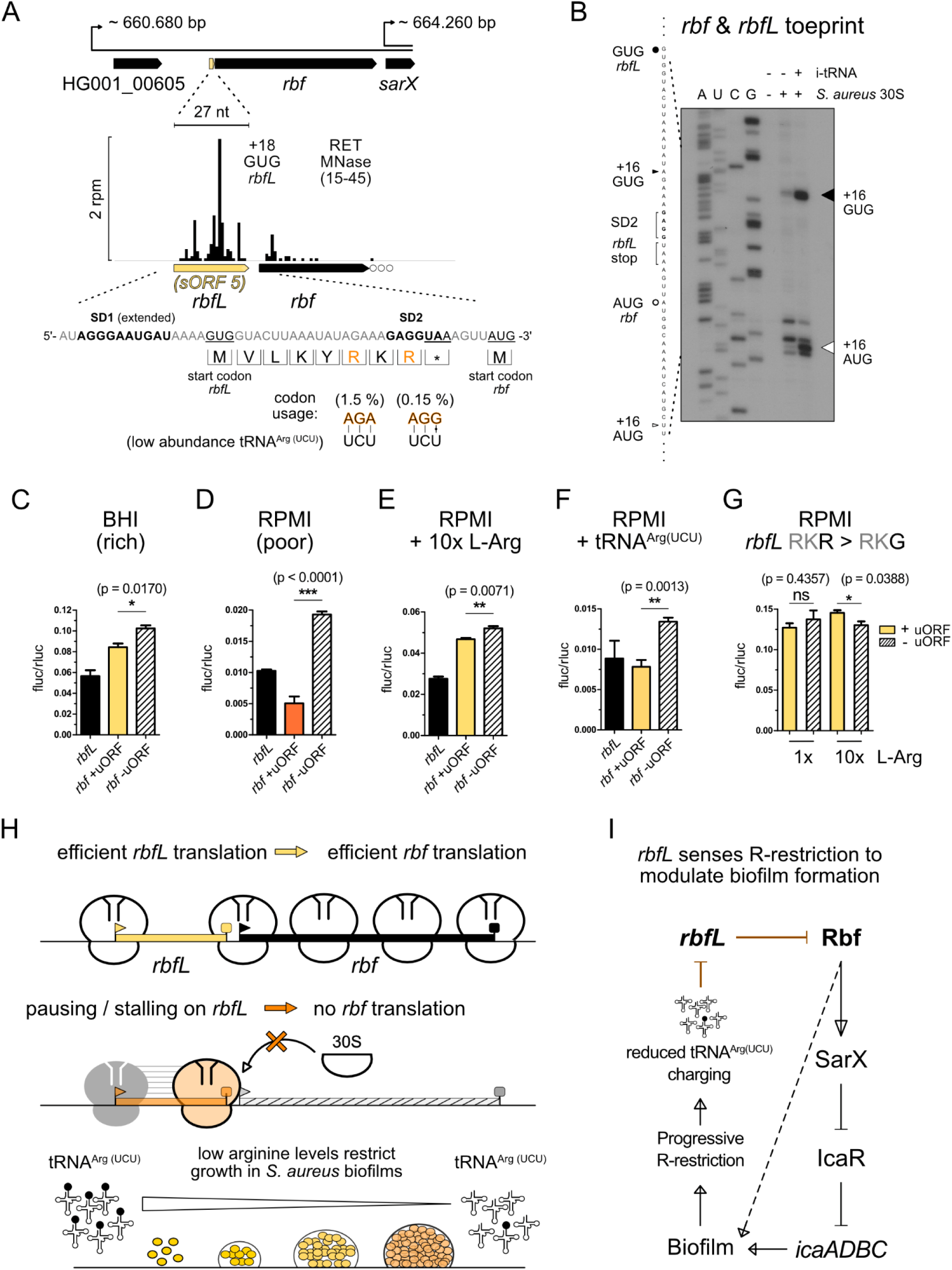
*rbfL* is a novel leader peptide, sensing arginine restriction and exerting translational control over a transcriptional regulator of biofilm formation. (A) Genomic organization of the *rbf* gene locus with its newly discovered leader peptide *rbfL* is depicted alongside a zoomed in MNase Ribo-Ret profile of corresponding ribosomal density peaks. The translation initiation sites of *rbfL* and *rbf* with their respective nucleotide sequence context are shown. SD motifs and start codons are highlighted bold and underlined, respectively. *rbfL* peptide translation is displayed, emphasizing decoding of a rare arginine codon (AGG). (B) Toeprint analysis demonstrating efficient 30S initiation complex formation at the start sites of *rbf* and *rbfL in vitro*. The corresponding sequencing lanes (A, U, C, G) and toeprint positions (+16) are indicated. (C, D, E, F, G) Results from *in vivo* dual luciferase assays measuring translation initiation efficiencies from *rbfL* and *rbf* reporter fusions in *S. aureus*. Variable culture conditions, such as growth media or L-Arg supplementation are indicated accordingly. Expression of *rbf* from its native sequence context in the presence of *rbfL* is highlighted in yellow, whereas *rbfL* start codon mutations are indicated by striped white bars. The strong regulatory effect of *rbfL* in RPMI is emphasized by the change from yellow to orange color for the *rbf* reporter fusion in panel D. Further reporter plasmid changes are indicated, corresponding to additional plasmid encoded tRNA^Arg(UCU)^ expression (F) or rare arginine AGG to GGG codon mutation (G), respectively. Measurements from Firefly luciferase reporter fusions were normalized by bi-cistronic Renilla luciferase expression. Mean and standard deviation from at least three biological replicates are shown. Two-tailed unpaired t test, ∗∗∗p < 0.001. (H) Graphical model of *rbf* translation regulation conferred by the *rbfL* leader peptide. (I) Schematic model linking a possible sensory role of *rbfL* translation to metabolic feedback control over *S. aureus* biofilm formation.

Although *rbfL* exhibited low expression during exponential planktonic growth, its discovery was enabled by the high sequencing coverage in our MNase data (Figure 6A). Ribo-Ret density mapped to a GUG start codon with an appropriately spaced **AGGGA**A**UGAU** extended SD motif and designated an eight amino acid small leader peptide (MVLKYRKR). Toeprinting confirmed efficient 30S initiation complex formation at the predicted *rbfL* GUG start codon, as well as at the downstream *rbf* start site (Figure 6B). Because the stop codon of *rbfL* is part of the SD motif of *rbf*, we hypothesize that *rbf* translation might be tightly coupled to efficient translation of its upstream leader peptide, *rbfL*. To test this hypothesis, we constructed dual luciferase reporter fusions, linking the *rbf* translation initiation region with Firefly luciferase and including its natural upstream sequence encompassing *rbfL*. As previously, Firefly luciferase was normalized by bi-cistronic Renilla luciferase expression. An additional construct fused to the eight amino acid *rbfL* CDS was used to monitor leader peptide translation. These constructs allowed us to assess the influence of *rbfL* on downstream *rbf* translation across conditions. In rich BHI medium, mutating *rbfL’s* GUG start-to a UAG stop codon only had a modest effect, resulting in a ∼10% increase in *rbf* translation efficiency (Figure 6C). However, under nutrient-poor conditions (RPMI medium), this effect of the leader peptide was markedly enhanced. Indeed, *rbfL* translation efficiency dropped 6-fold, while *rbf* expression was strongly *rbfL*-dependent, increasing ∼4-fold in the absence of the leader peptide (Figure 6D). Together, these findings suggest that nutrient availability modulates *rbf* translation efficiency in an *rbfL*-dependent manner.

Upon inspecting the *rbfL* coding sequence, we noted that one of its two arginine codons was a particularly rare AGG codon (0.15% usage frequency). Both arginine codons (AGA and AGG) are decoded by the relatively scarce tRNA^Arg(UCU)^. We hypothesized that limited availability of the charged tRNA^Arg(UCU)^ could constrain *rbfL* translation, thereby modulating downstream *rbf* expression. To test this, we repeated the dual-luciferase reporter assays in RPMI medium supplemented with 10-fold excess of L-Arg (0.2 → 2.2 g/L). Strikingly, this supplementation increased *rbfL* translation efficiency three-fold (relative to the Renilla luciferase) and fully relieved its repressing effect on *rbf* translation (Figure 6E). To further explore the role of tRNA^Arg(UCU)^ availability, we introduced an additional plasmid-borne copy of the *argU* gene (encoding tRNA^Arg(UCU)^), expressed from its natural promoter. This partially alleviated *rbfL*-dependent translational repression, as *rbf* expression in the absence of *rbfL* was only 1.7-fold higher compared to a 4-fold increase without *argU* (Figure 6F). Finally, to directly assess the role of the rare arginine codon, we substituted the AGG codon with a glycine codon (GGG), ensuring that this mutation did not disrupt SD-aSD base pairing with the overlapping *rbf* RBS (Figure 6A). Consistent with our hypothesis, this single-nucleotide change fully abolished *rbfL*-dependent translational repression and rendered *rbf* expression insensitive to L-Arg supplementation (Figure 6G). Collectively, these findings support a mechanism of RBS occlusion, in which slow decoding or ribosome stalling on the *rbfL* leader peptide impedes translation initiation at the downstream *rbf* start codon (Figure 6H). Our data suggest that arginine availability is the primary cue sensed by *rbfL* through the charging status of tRNA^Arg(UCU)^.

## Discussion

Regulation of translation initiation is central to bacterial adaptation, but species-specific mechanisms remain poorly understood. In this study, we investigated translation initiation and its regulation in *Staphylococcus aureus* by high-resolution initiation profiling with Ribo-Ret. While RNase 1 has previously been used for standard Ribo-seq in *Listeria* and *Salmonella* (47), our work provides the first high resolution, genome-wide dataset of translation initiation sites in bacteria. Although Ribo-Ret has gained popularity predominantly for sORF discovery (48, 58–61), we exploited its high precision obtained with RNase 1 to explore translation initiation mechanisms at single-nucleotide resolution. By combining these profiling data with cryo-EM analysis using an endogenous *S. aureus* mRNA, we uncover the structural basis for species-dependent start codon selection, revealing that *S. aureus* 70S ribosomes accommodate a shifted and extended SD-aSD duplex with additional base pairing as compared to *E. coli*. This structural stabilization rationalizes previously noticed differences in preferred spacing between the SD and start codon across bacteria (25, 62) and provides a mechanistic explanation for divergent start codon selection in *S. aureus*. In bacteria, the SD-aSD helix provides an anchoring point for the mRNA during its accommodation on the ribosome. An optimal distance between the SD-aSD helix and the start codon ensures proper positioning of the start codon in the ribosomal P-site, facilitating its recognition by the initiator tRNA. In *S. aureus*, the extended SD-aSD helix influences how start codons are presented in the ribosomal P-site (Figure 3). The rigidity introduced by the additional start codon proximal interactions limits mRNA flexibility. Together with the shifted position of the extended SD-aSD helix on the ribosomal platform, they contribute to the specific optimal spacer length that is required to correctly position the start codon in the P-site (Figure 2). In *E. coli*, a shorter SD-aSD helix with slightly different position on the platform, requires shorter spacing for optimal localization of the start codon. In the absence of alternative, competing start codons, suboptimal spacer length incurs only a modest energetic cost, lowering initiation efficiency (22). However, when two potential start codons are present in the same RBS, the optimally spaced start codon will be preferred over the suboptimally positioned alternative. Strikingly, in these situations, the distinct spacer preferences observed among different bacterial species can lead to species-dependent decoding outcomes. This is demonstrated by our characterization of start codon selection in the context of *aur* and *rlmB* mRNAs, where *S. aureus* and *E. coli* initiate translation of different open reading frames (Figure 2C and D). The potential for extended SD-aSD base-pairing implies that a comparatively larger portion of the 16S rRNA is accessible for interaction with the mRNA, thereby allowing complementary sequences beyond canonical motifs to function as effective SD elements. In species capable of forming extended SD-aSD helices, the definition of SD motifs becomes less restrictive, encompassing a broader range of sequences. For example, in the *rarA* and *proP* mRNAs, such atypical alignments may underlie the accurate and efficient start codon recognition observed in *S. aureus*, but not in *E. coli* 30S initiation complexes (Figure S4). While we do not define the full spectrum of these interactions in *S. aureus*, our data clearly show that extended SD-aSD pairing is a key determinant of start codon selection and contributes to evolutionary distinct patterns of ORF recognition. These findings underscore the potential to engineer mRNAs that produce distinct proteins depending on the bacterial species that translate them. This concept has broad implications for biomedical and biotechnological applications, including precise control of protein expression, species-specific translational switches in synthetic biology, and the targeted design of pathogen-specific antibacterial strategies.

Given the conserved 3’-end sequence of the 16S rRNA between *S. aureus* and *E. coli*, other features must account for the mechanistic divergence that enables extended SD-aSD formation and distinct start codon selection (Figures 2 and 3). One notable difference is the presence of ribosomal protein bS21 near the SD-aSD helix in *E. coli*, which would sterically hinder the extended interaction observed in *S. aureus*. In Firmicutes, bS21 is relatively shorter and lacks the C-terminal residues needed to anchor it to the 16S rRNA (63), consistent with the absence of a corresponding EM-density in our structure (Figure 3). Notably, bS21 has previously been linked to evolutionary translation initiation adaptations (17). Another important difference is the absence of ribosome-anchored protein bS1 in *S. aureus* (64). In *E. coli* bS1 is essential and bound at the ribosomal platform to assist translation initiation by lowering stability of structured ribosomal binding sites (10). Further studies are required to clarify the roles of bS21 and bS1 in modulating translation initiation in *S. aureus*. Which evolutionary pressure might have driven the selection of extended SD-aSD base pairing? We hypothesize that this adaptation could be linked to genome composition, specifically, the high AT content in species like *S. aureus*, and the need to adapt their translation initiation strategy accordingly. In *E. coli*, which has a more balanced GC content, ribosomes predominantly rely on an unstructured region near the start codon for recruitment, and the SD-aSD pairing is not critically required (17). In contrast, AT-rich genomes, like those of many Firmicutes, may rely more strongly on stable SD-aSD interactions. This could be due to the generally lower mRNA secondary structure in these organisms (65), making it more difficult to define initiation sites by their unstructured nature.

Our findings helped define which RBS sequences strongly promote translation initiation, hence identifying sORF candidates with greater confidence and enabling the discovery of novel non-canonical start codons. In *S. aureus*, translation attenuation via non-canonical start codons has not previously been described. This mechanism typically keeps gene expression low through inefficient decoding in the ribosomal P-site, a process modulated by IF3 (66, 67), which itself uses an AUA start codon in *S. aureus*, likely enabling auto-regulation as seen in *E. coli* (68). Using Ribo-Ret, we identified two additional conserved non-canonical start codons limiting initiation efficiency of *S. aureus cvfC1* and *ccpN* mRNAs. The strong conservation of *ccpN*’s AUA start codon may be linked to CcpN’s function as a transcriptional repressor of gluconeogenesis (57, 69). Strikingly, we observed that *ccpN’s* initiation efficiency varies in a growth media-dependent manner. Under nutrient-rich conditions, translation initiation increases, hinting at a potential metabolic feedback control mechanism. Efficient *ccpN* translation could repress gluconeogenesis when nutrients are abundant, and inefficient translation would relieve this repression under starvation.

We have identified 43 novel sORFs and validated several initiation sites using toeprinting and reporter fusions (Supplementary Table S1 and Figure 5E, respectively). Especially, the short uORFs are challenging to detect with standard methods like mass spectrometry or regular Ribo-seq. In these cases, Ribo-Ret’s strong enrichment at start sites determined at high resolution was critical to enable their identification. While our focus was primarily on regulatory uORFs, several of the newly-identified sORFs (e.g., *sORF25*, *26*, *28*, *30*, *36* and *45*) are predicted to adopt hydrophobic transmembrane helices, suggesting possible functional roles. Furthermore, *sORF36* and others (*sORF6*, *26*, *30*, *34*, *35* and *40*) overlap with sRNA annotations, suggesting that some of them may represent dual-function regulatory RNAs (70), opening future prospects for further functional characterization. Short uORFs are often associated with regulatory functions. In bacteria, tRNA charging levels can reflect nutrient availability, and leader peptides can act as sensors of amino acid scarcity by causing ribosome pausing, thereby regulating downstream gene expression. In *S. aureus*, a leucine-rich leader peptide was described to regulate isoleucine biosynthesis by promoting transcription antitermination under branched-chain amino acid limitation (53). Here, we describe a different mechanism, involving leader peptide-dependent regulation of translation initiation. Strikingly, we show that a novel leader peptide, *rbfL*, senses arginine limitation through the presence of a rare arginine codon, which is decoded by the low-abundance tRNA^Arg(UCU)^. When arginine is scarce, ribosomes pause or stall on *rbfL*, blocking the ribosome binding site of the downstream gene *rbf*, and preventing its translation initiation. Given that Rbf is an important transcription factor promoting biofilm formation, this regulation likely restricts excess of biofilm formation under arginine-limiting conditions (Figure 6I). Importantly, pronounced arginine limitation was recently shown to restrict protein synthesis and growth in *S. aureus* biofilms, contributing to antibiotic tolerance (71). Our finding also aligns with previous studies linking arginine metabolism and biofilm formation in *S. aureus*, including the role of the sRNA RsaE in promoting biofilm formation (72) and repressing arginine catabolism (73). The arginine-sensitive control of biofilm formation via r*bfL* further strengthens the connection between arginine metabolism and biofilms in *S. aureus*. While our study focused on *rbfL*, our data suggests the existence of other regulatory leader peptides, offering exciting directions for future research.

## Methods

**Resource table**

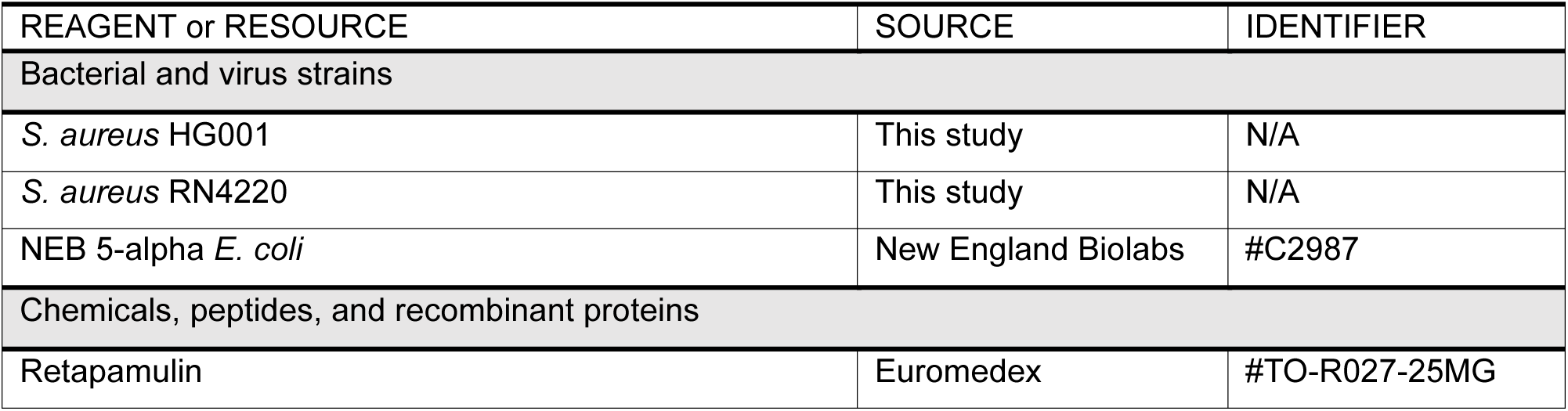
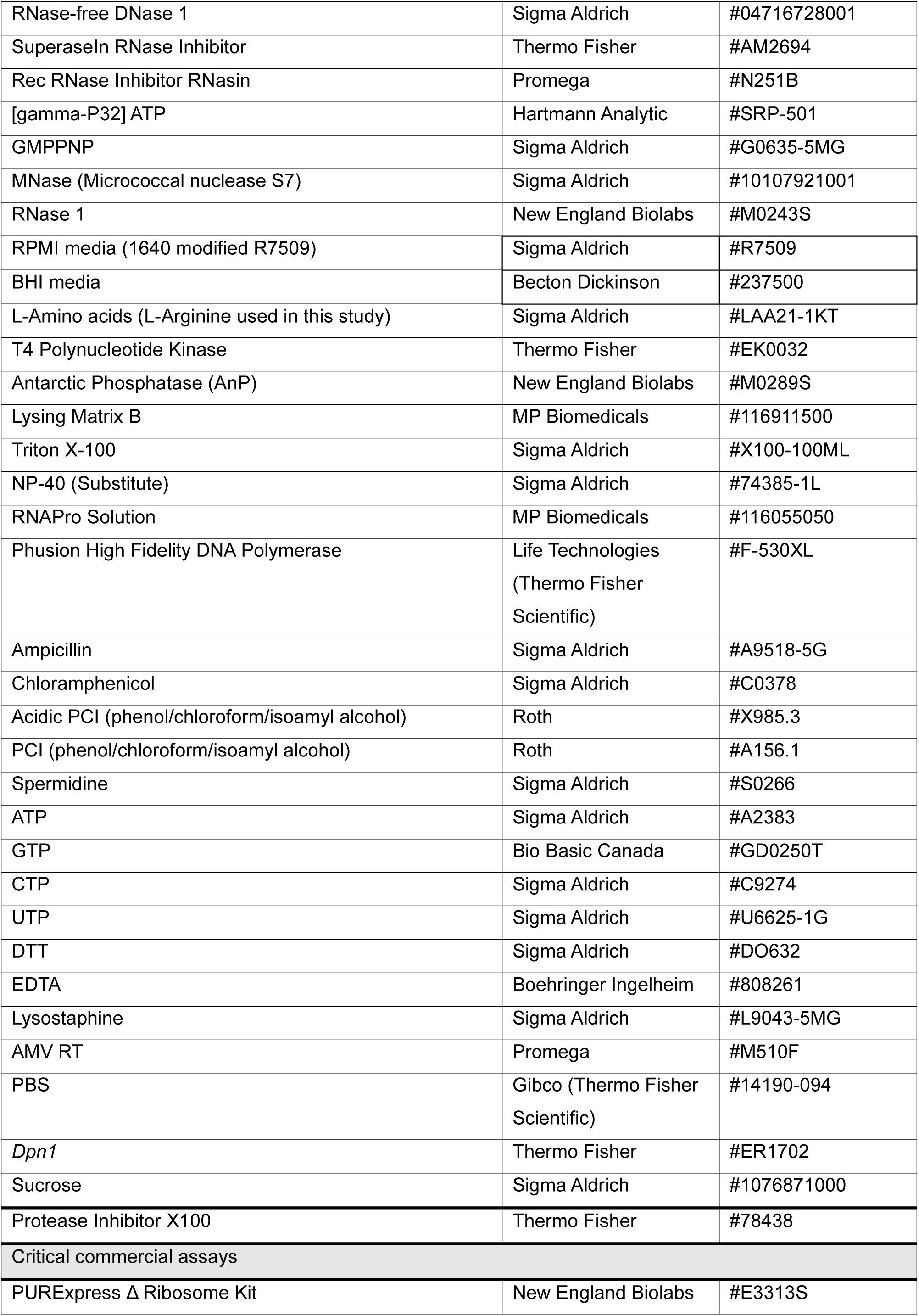
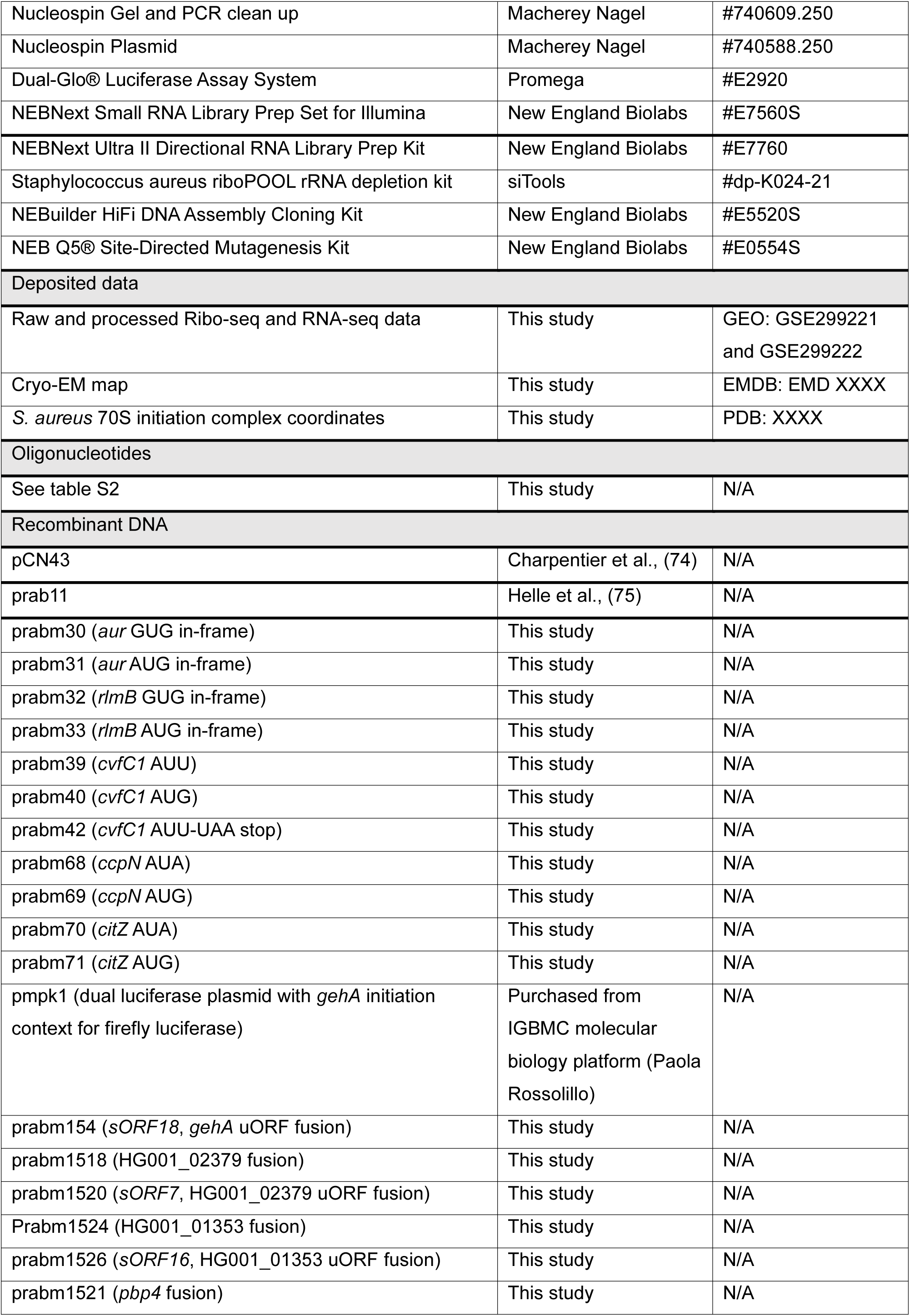
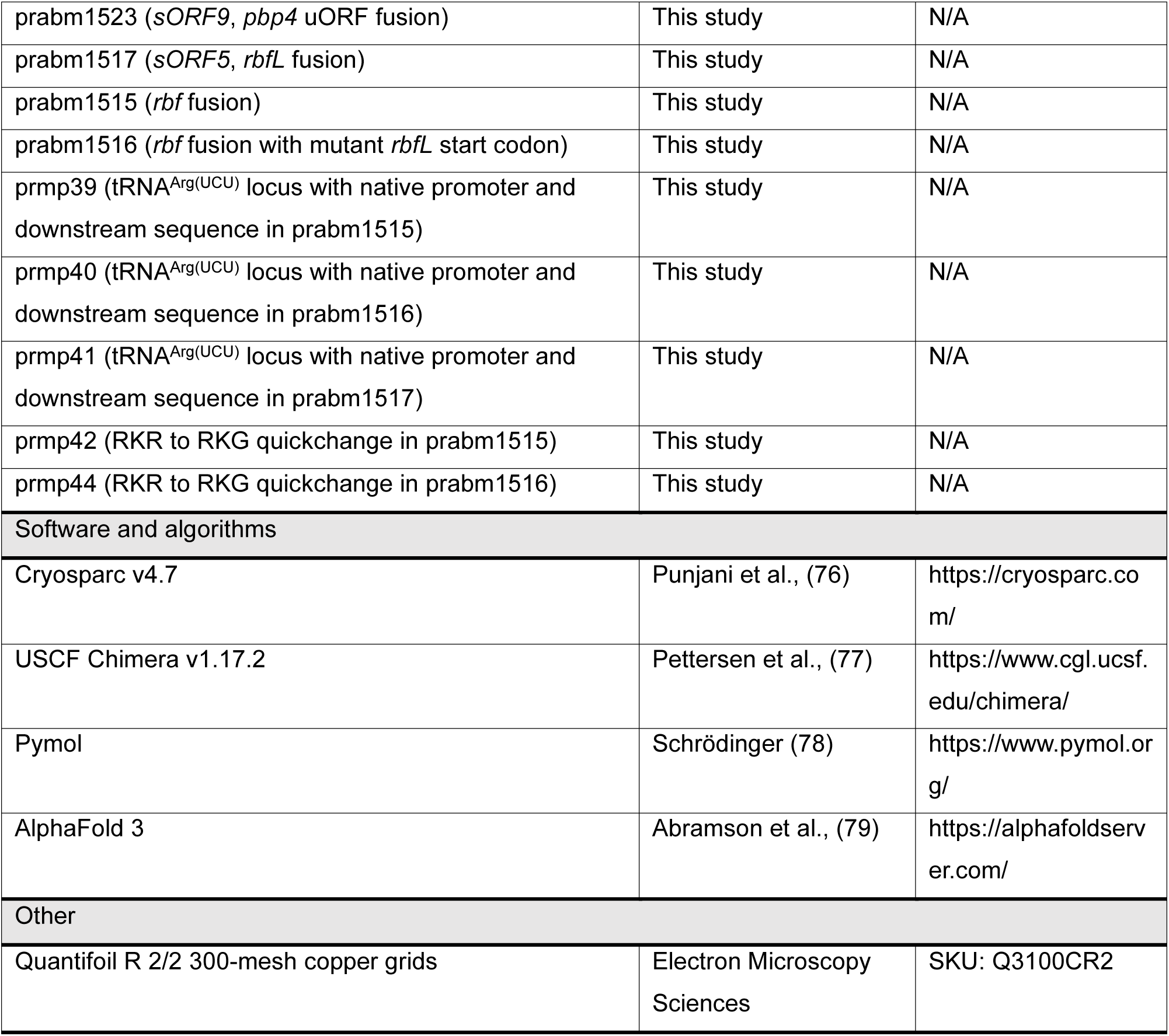

### Ribo-seq experiments

The Ribo-seq and Ribo-Ret experiments were performed as previously described (44), largely following the procedure for Ribo-Ret in *E. coli* (48) with several adjustments based on published recommendations (46, 61, 80, 81) and on our own optimization for *S. aureus* (44). For this study, few key steps of the protocol were changed. *S. aureus* HG001 was plated and grown at 37°C o/n on BHI agar plates. Single colonies were used to inoculate 3 ml of BHI and precultures were grown, shaking at 180 rpm, o/n at 37°C. The following day, precultures were diluted to an optical density A_600_ (OD) of 0.05 in 50 mL of BHI in 250 ml flasks. Bacterial cultures were grown, shaking at 180 rpm at 37°C, to an OD of 3.5 and then briefly (5 min) treated with 25 μl of 25 mg/ml Retapamulin (12.5 μg/ml f.c.) for Ribo-Ret. From then, Ret-treated and non-treated samples were processed identically for cell harvest and lysis.

Cultures were subjected to rapid cooling by swirling in an ice-bath for 3 min, before 40 ml of each sample (split into two 50 ml falcon tubes) were harvested by centrifugation at 2,100 g for 8 min. Another 10 ml of each culture were flash frozen in liquid nitrogen and reserved for total RNA sequencing, as previously described (44). For Ribo-seq and Ribo-Ret, cell pellets were resuspended in 550 μl of pre-chilled lysis buffer A (20 mM Tris-HCl pH 8, 50 mM MgCl_2_, 100 mM NH_4_Cl, 5 mM CaCl_2_, 0.4% Triton X-100, 0.1% NP-40, 100 U/mL DNase 1, 1 mM GMPPNP). Mechanical lysis was performed by two rounds of ‘fastprep’ bead-beating at 6 m/s for 40 seconds in 2 mL fastprep tubes containing Lysing Matrix B. Subsequently, samples were centrifuged at 16 000 g for 5 min at 4°C and supernatants were carefully retrieved. Measurement of absorbance at 260 nm in a Nanodrop spectrophotometer served to adjust nuclease concentration for subsequent digestion by either MNase or RNase 1. For MNase digestion, 750 U of MNase and 50 U of SUPERase-In (does not inhibit MNase) were added per 40 absorbance units (AU). After 1 h shaking at 1400 rpm and 25°C, the reactions were quenched by addition of 5 mM EGTA (f.c.). For the alternative digestion by RNase 1, 250 U of RNase 1 were added per 40 AU of sample concentration. The digestion was performed for 20 min shaking at 800 rpm and 37°C, before it was stopped by addition of 100 U of SUPERase-In. All samples were kept strictly on ice prior to loading onto a 5-50% sucrose gradient for isolation of 70S monosome peaks. Ribosome protected fragments (RPFs) were extracted using acidic hot phenol and size selected on a 15% polyacrylamide-8 M Urea gel (PAGE) as previously described (44) with only minor changes. Namely, RPFs were gel eluted via passive elution in RNA elution buffer (0.5 M NH_4_Ac pH 6.5, 1 mM EDTA, 0.1% SDS) o/n at 4°C shaking at 700 rpm prior to acidic PCI (phenol/chloroform/isoamyl alcohol, 25/24/1) extraction and ethanol RNA precipitation.

### Library preparation and sequencing analysis

All samples were subjected to ribosomal RNA depletion employing a commercially available rRNA depletion kit according to the manufacturer’s protocol. Prior to library preparation, RPFs were de-phosphorylated by the use of Antarctic phosphatase and subsequently phosphorylated with T4 polynucleotide kinase. RPF libraries were then prepared using the NEBNext Small RNA Library Prep Set from Illumina, following the kit’s instructions. For the associated total RNA sequencing libraries, a different commercially available kit was employed (NEBNext Ultra II Directional RNA Library Prep Kit). After single-end sequencing on an Illumina NGS instrument, demultiplexed sequencing data in FASTQ format were processed as previously described (44), following published and publicly available workflows (48, 80). To assess ribosome enrichment at start codons globally and to compare resolution of translation initiation peaks between MNase and RNase 1, a global average gene analysis was performed as previously described (48).

### sORF mapping

For the discovery of novel, unannotated small ORFs (sORFs) we followed a similar strategy as described by Meydan et. al (48). After *in silico* translation, all potential ORFs with AUG, GUG or UUG start codons were assigned to Ribo-Ret peaks from duplicate MNase datasets, where 3’ ends of reads mapped within a window of +15 to +21 nt, where e.g., A of AUG designates position +1. Because of higher absolute sequencing coverage (> 25 million vs. approx. 10 million mapped reads per sample), we initially utilized the MNase datasets to search for densities corresponding to novel unannotated sORFs, and later visually inspected all candidate sORFs using the densities of the high-resolution RNase 1 datasets. For this analysis, we took into account reads between 32-45 nt in length, additionally enriching translation initiation peaks. We then proceeded to filter the dataframe containing all potential unannotated ORFs based on differential rpm expression thresholds (see Supplemental Scheme S1). We considered all intergenic, unannotated ORFs but also ‘hypothetical protein’ annotations below 100 aa for this analysis and later focused on sORFs below 50 aa in length. Based on SD motif presence (’GGGGG’, ‘GGGG’, ‘GGAGG’, ‘GAGG’, ‘GGAG’, or ‘AGGA’) 16-4 nts upstream of start codons, we considered ORFs with RET peak values above 0.1 rpm for visual inspection, but were more stringent with a minimum of 1 rpm in both duplicate experiments for ORFs without predicted SD-motif. Because of low, but consistent background levels of Ribo-Ret density within annotated coding sequences globally, presumably due to high levels of pervasive translation, we did not focus our analysis on potential internal initiation sites and excluded them from this mapping approach. Rather, we visually inspected the resulting 944 candidate intergenic sORF initiation sites individually and compared Ribo-Ret densities with the equivalent RNase 1 data. We narrowed down our selection based on mutual expression in the MNase and RNase 1 datasets and constructed a high confidence sORF list. Notably, the majority of final considered densities corresponded to sORFs which had strong SD-motif predictions and which had RNase 1 density peaks mapping at appropriate distances to the predicted start codons. After final genome-wide data examination in an integrative genome viewer (IGV), 5 additional sORFs were added manually, which had been missed computationally, either due to alternative annotations (e.g., putative phage protein) or because their densities had mapped diffusely, while other criteria such as strong SD-motif and appropriate aligned spacing supported sORF presence. The final curated sORF list is provided in Supplementary table S1.

### Aligned spacing analysis

The comparatively small genome size of *S. aureus* had allowed us to visually inspect all translation initiation sites on a genome browser, individually assessing precise start codon selection, resulting in our latest, improved *S. aureus* HG001 annotation. To study differences in start codon selection and spacing preferences, we calculated the average aligned spacing of *S. aureus* HG001 and the *E. coli* K12 reference genome. The first 20 nt upstream of all annotated start codons were extracted and a motif search was performed considering all instances of ‘GAGG’, ‘GGAGG’ and ‘GGAG’. Exclusively strong, core SD-motifs were considered to avoid ambiguous SD-aSD pairing during comparison. For the same reason, sequences with multiple potential SD-motifs were excluded from the analysis, unless the motifs fully overlapped. In few cases of multiple identical sequences extracted, only one instance was considered to avoid confounding effects by duplicate annotations. The nucleotide sequence distance to the start codon was finally counted from the end of the three defined motifs for all upstream sequences identified. To align spacing at the 3’ end of the GGAGG core motif, a one nucleotide spacing count was subtracted in cases of GGAG motifs. The absolute counts of identified motifs were plotted according to their computationally determined aligned spacing (Figure 2A).

### Extended start site analysis

The 20 nt sequences upstream the start codon, extracted for the aligned spacing analysis, were used to generate a sequence logo, utilizing the Logomaker Python package. Following our biochemical analysis of extended SD-aSD base pairing, these sequences were re-analysed for SD-motif (SD+) and extended SD-motif (SD++) presence. To check for the general presence of SD motifs, the first 16 nt (−20 to −5) were searched for any occurrences of GAGG, GGAG, GGGG, GGAGG, AAGG, AGGA, AGGT or GGGT sequences. For all sequences where the general presence of an SD-motif was identified, we also searched the full 20 nt upstream sequence for occurrences of AGGNAT, AGGNGT, GGGNAT or GGGNGT, where ‘N’ denotes any nucleotide, to predict the formation of an extended SD-aSD helix. Finally, to analyse the start codon usage, the first three nucleotides of the CDS were extracted and different start codon usage counted.

### Nucleotide sequence conservation analysis

Nucleotide sequence conservation of individual genes has been regularly assessed using curated genomic data publicly available through the SEED database (82). *S. aureus* genes of interest were first searched for homologs in the database. Their respective coding sequences and upstream nucleotide sequences were extracted, and potential SD motifs and translation initiation sites individually assessed.

### *In vitro* transcription

*In vitro* transcriptions were routinely performed on template PCR products harbouring T7 promoter sequence overhangs. Following standard PCR procedures with Phusion DNA polymerase, PCR products were assessed on 1% agarose gels and purified using a commercially available clean-up kit (NucleoSpin Gel and PCR Clean-up, Macherey Nagel). *In vitro* transcription assays were performed in 400 μl reaction volumes, with a minimum of 1 µg of purified PCR product. Typical reactions contained 4 mM NTPs (final concentration) each, 0.2 mM Spermidine, 5 mM DTT, 4 μl of RNase inhibitor (RNasin) and 8 μl of home-made T7 RNA polymerase. Reactions were adjusted to 1x reaction conditions making use of a 10x *in vitro* transcription buffer (0.4 M Tris-HCl pH=8 at 37°C, 150 mM MgCl_2,_ 0.5 M NaCl). *In vitro* transcriptions were performed for 3 h at 37°C. Subsequently, 40 μl of DNase 1 buffer 10x and 10 μl of DNase 1 were added and template PCRs were digested for 1 h at 37°C. DNase digestion was stopped by addition of 0.5 M EDTA and the resulting RNA was purified by extraction with acidic PCI (phenol/chloroform/isoamyl alcohol, 25/24/1) solution followed by ethanol precipitation. RNA products were subsequently purified on a preparative 6% PAGE and excised from the gel by UV-shadowing. Correct product sizes were gel eluted via passive elution in RNA elution buffer (0.5 M NH_4_Ac pH 6.5, 1 mM EDTA, 0.1% SDS) o/n at 4°C shaking at 700 rpm. A final acidic PCI phenol/chloroform extraction was followed by ethanol precipitation. Purity and quantity were routinely assessed by Nanodrop spectrophotometry.

Notably, the initial PCR reactions for T7 template generation were frequently performed with sequence-modified ssDNA oligo sequences as forward primers to introduce the desired nucleotide changes in the PCR template. This has been done, for example, to generate sequence variants for the analysis of the mutant *spa and aur* RBS sequence contexts, analysed by toeprinting.

### 30S toeprinting analysis

Toeprinting (83) of *S. aureus* or *E. coli* 30S initiation complexes was performed as previously described (84). Ternary complexes were formed with *in vitro* transcribed mRNA, uncharged initiator tRNA^fMet^, and purified 30S subunits as follows: 1 pmol of mRNA was hybridized with 2-4 pmol of ^32^P-radiolabeled ssDNA toeprinting oligo in a 9 μl reaction volume, briefly denatured for 1 min at 90°C and refolded by incubation for 1 min on ice and then at room temperature for 5 min. Subsequently, 1 μl of toeprint buffer 10x (100 mM MgCl_2_, 200 mM Tris–HCl of pH 7.5, 600 mM NH_4_Cl and 10 mM DTT) was added. Before use, purified 30S were adjusted to 2 µM concentration in toeprinting buffer 1x and pre-incubated at 37°C for 15 min. Subsequently, 2 µl of mRNA/oligo mix, 2 µl of 30S and 1 µl of toeprint buffer 10x were added in a final reaction volume of 10 μl. After 10 min incubation at 37°C, 1 µl of 20 μM *E. coli* deacylated initiator tRNA^fMet^ was added and samples were incubated for another 5 min at 37°C. For primer extension, a master mix was prepared so that addition of 2 μl would result in final concentrations of 0.2 mM dNTPs, 1x AMV RT buffer and 1 U/µl of AMV RT. Primer extensions were performed for 30 min at 37°C. Subsequently, RNA templates were hydrolysed by addition of 3 μl of 3 M KOH as well as 20 μl of buffer X (50 mM Tris-HCl pH 8.0, 0.5% SDS and 7.5 mM EDTA) and incubation for 2 min at 90°C. Subsequently, 6 μl of 3 M acetic acid were added to neutralize the reactions. In the final step, cDNA was recovered by Phenol-Chloroform extraction and ethanol precipitation with addition of sodium-acetate (0.3 M f.c.). cDNA pellets were resuspended in urea loading dye, radioactivity counts were normalized and equal amounts resolved on a denaturing 10% PAGE. Gels were exposed to Fuji X-ray films and developed using an Optimax X-ray film processor. Sequencing lanes were generated as previously described (84).

### 70S toeprinting analysis

For toeprinting of 70S complexes during *in vitro* translation, a specialized kit was used *(PURExpress delta ribosome kit, NEB*). Toeprinting of *in vitro* translation reactions was performed as previously described (21). In brief, following the manufacturer’s protocol, a typical reaction contained 2.5 μl of solution A, 0.75 μl of Factor-mix, 1 pmol of gel-purified mRNA template and 0.5 μl of RNase inhibitor (RNasin) in 6.25 μl final reaction volumes. Then, 8 pmol of purified *E. coli* or *S. aureus* 70S ribosomes were supplied. To stall 70S complexes with Retapamulin its concentration was adjusted to 50 μM in the final reaction volumes. *In vitro* translations were performed for 30 min at 37°C prior to addition of 1-2 pmol of ^32^P-radiolabeled ssDNA toeprinting oligos. Subsequently, the reactions were supplied with 2 μl of a primer extension mix, providing the four dNTPs at 0.4 mM f.c. each and 2 U of AMV RT per reaction, adjusted to 1x reaction conditions with commercial 5x AMV RT buffer. Primer extensions were performed directly in the toeprint reactions for 30 min at 37°C. Following cDNA Phenol chloroform extraction, toeprint reactions were resolved as described for 30S toeprinting.

### Reporter plasmid construction

Reporter plasmids were routinely cloned making use of NEBuilder HiFi DNA Assembly to introduce various inserts into a published ‘prab’ vector plasmid backbone (75), carrying constitutively expressed ampicillin and chloramphenicol resistance genes. We had initially introduced a bi-cistronic Renilla- and Firefly reporter cassette, constitutively co-transcribed from a P_Blaz_ promoter sequence into a different vector backbone (pCN43) (74). However, because of the pcN43 encoded erythromycin resistance, we chose to transfer the cassette with its promoter, into the prab vector series with the chloramphenicol resistance. The rationale was to avoid any artificially introduced modifications on the ribosome, which could potentially interfere with the interpretation of our *in vivo* assays on bacterial start codon selection or ribosome stalling on leader peptides. Inserts for reporter fusions were regularly obtained amplifying PCR products with plasmid-complementary overhangs from the upstream sequences and translation initiation sites of genes of interest from HG001 gDNA. These inserts were cloned in-between the bi-cistronic context of the upstream Renilla- and downstream Firefly luciferase sequences, creating translational reporter fusions with the CDS of the latter.

Assemblies were generally performed following the manufacturer’s protocol, but scaled to a total reaction volume of 4 μl. Both vector backbone and insert PCRs were generally assessed on 1% agarose gels, and spin-column kit purified following the manufacturer’s protocol. Typically, 50 fmol of purified vector backbone PCR were mixed with 2-5-fold molar excess of purified insert PCRs for assembly with 2x assembly master-mix. Following 1h incubation at 50°C, the reactions were directly transformed into 50 μl of NEB 5-alpha or home-made competent top10 *E. coli* cells, by standard heat-shock procedures.

For simpler nucleotide sequence changes, either the Q5 site-directed mutagenesis kit or an alternative ‘quick-change’ protocol was employed. Q5 site-directed mutagenesis was performed according to the manufacturer’s protocol. For the alternative quick-change protocol, complementary 35-45 nt ssDNA oligos were purchased with matching plasmid sequence context, except for 1-2 nt mismatches at the centre of the oligos, designating desired plasmid sequence changes. Between 50 and 200 ng of vector plasmid served as template for amplification by Phusion PCR. Subsequently, 1 μl of *Dpn*I was used to digest template plasmid vector for 1 h at 37°C, prior to heat-shock transformation.

Plasmids were routinely purified by miniprep utilizing a commercially available kit. All required DNA oligonucleotides were purchased from Integrated DNA Technologies (IDT). Every cloned reporter construct was individually verified by Sanger sequencing (Eurofins genomics).

### *S. aureus* reporter plasmid transformation

Due to their inherent ease of transformation, *S. aureus* RN4220 cells were generally employed for *in vivo* expression of various reporter plasmids. Competent RN4220 cells were prepared and transformed as follows: RN4220 glycerol stock was initially inoculated into 3 ml of BHI for a preculture, grown at 37°C and shaking at 180 rpm o/n. The following day, a fresh dilution to an OD of 0.05 in 50 ml BHI was grown under equivalent conditions until reaching OD 0.6. At this stage, cells were split into two 50 ml falcon tubes and pelleted in a pre-chilled centrifuge at 4°C for 10 min at 2,100 g. Pellets were washed twice by resuspension in 10 ml of 0.5 M sucrose and subsequent centrifugation. After discarding the supernatants, cell pellets were resuspended in 300 μl 0.5 M sucrose each. Then, 100 μl of cells were mixed with 10 μl of plasmid at approximately 500 ng/μl concentration and directly subjected to electroporation. Subsequently, 900 μl of BHI were added, and the cells were transferred to 1.5 ml tubes and left to recover at 37°C 2 h shaking at 180 rpm. After recovery, transformed cells were pelleted in a microcentrifuge, resuspended in 100 μl BHI and plated on BHI agar plates containing 10 μg/ml chloramphenicol (Cm) as a plasmid selection marker.

### *In vivo* dual luciferase reporter assays

Dual luciferase reporter plasmids were routinely transformed into RN4220 competent *S. aureus* cells, selected via the plasmid-conferred chloramphenicol resistance and stored as glycerol stocks at −80°C. To perform the dual luciferase assays, these stocks were inoculated into pre-cultures in 3 ml of BHI media containing 10 μg/ml Cm, and grown at 37°C, shaking at 180 rpm o/n. For each biological replicate, a separate pre-culture was inoculated. The following day, pre-cultures were diluted 1:100 in 5 ml of fresh BHI or RPMI media, containing 10 μg/ml Cm, in 50 ml falcon tubes and grown at 37°C shaking at 180 rpm. For rich BHI media, cultures were generally grown for 4 h, reaching mid to late exponential growth phase, whereas cultures in RPMI were grown for up to 7 h. Because of the difference in bacterial growth, the lysis procedure slightly differed. For BHI cultures, 50 μl were directly transferred into 96-well plates where they were added to 50 μl of PBS solution containing 5 μg of Lysostaphin. Because of the lower optical densities obtained from 7 h of growth in poor RPMI media, 2 ml of these cultures were pelleted in a microcentrifuge and directly resuspended in the equivalent 50 μl of PBS solution containing 5 μg of Lysostaphin, which were then transferred into the 96-well plate instead. *S. aureus* cell lysis was achieved by subsequent incubation for 30 min at 37°C. Subsequently, 25 μl of lysate were transferred into an adjacent well, where the dual luciferase assay was performed. First 25 μl of Promega Dual-Glo firefly luciferase substrate was added and reactions were incubated for 10 min at room temperature. Subsequently, relative light units (RLU) were measured in a Promega GloMax luminometer. According to the manufacturer’s protocol, Firefly luciferase was then quenched by the addition of 25 μl of Renilla luciferase substrate reaction mixture (Dual-Glo Stop & Glo) and incubated for another 10 min. Finally, a second measurement of relative light units (RLU) in the luminometer, served to normalize the previously obtained Firefly signal intensities. For all reactions, a negative control was included from BHI or RPMI media assayed in parallel, whose low background signals were subtracted from sample datapoints before normalization.

Because the constitutive P_Blaz_ promoter is also active in *E. coli,* the initially transformed home-made competent top10 *E. coli* cells used for cloning could also be used to assay the transformed reporter plasmids *in vivo*. The respective protocol was largely consistent with the dual-luciferase assays performed in *S. aureus* and only slightly differed in terms of growth conditions and lysis. In detail, plasmid-containing glycerol stocks were inoculated in 3 ml pre-cultures of LB media containing 100 µg/mL ampicillin and grown at 37°C, shaking at 160 rpm o/n. The pre-cultures were diluted 1:100 in 5 ml fresh LB (Amp) and grown in 50 ml falcon tubes at 37°C, shaking at 160 rpm for approximately 4-5 h. Next, 2 ml of each culture were pelleted in a microcentrifuge and resuspended in 600 μl of PBS. For lysis, these resuspensions were transferred into 2 mL fastprep tubes containing Lysing Matrix B and subjected to two rounds of fastprep at the recommended *E. coli* protocol for mechanical lysis by the MP Biomedicals fastprep system. Subsequently, the tubes were spun down at 13 000 rpm in a microcentrifuge and 25 µl were directly transferred into 96-well plates for dual luciferase assay as described for *S. aureus* cell lysates.

### *S. aureus* ribosome purification

*Staphylococcus aureus* 70S ribosomes were purified as previously described (64). Briefly, fresh o/n precultures were diluted to an OD_600_ of 0.05 in two liters of BHI media and grown at 37°C shaking at 180 rpm to an OD_600_ of 1.0, reaching logarithmic growth phase. Cells were harvested by centrifugation at 2100 g at 4°C and washed twice with cold buffer A (20 mM HEPES-KOH pH 7.5, 100 mM NH_4_Cl, 21 mM Mg(CH₃COO)_2_,1 mM DTT). Approximately 1.5 g of bacterial cell pellet was resuspended in 10 ml of buffer A, supplemented with 0.75 mg of lysostaphin, 150 U of DNase 1, 24 μl of 0.5 M DTT as well as 50 μl of Protease Inhibitor X100 and then incubated at 37°C for 45 minutes. Cell lysate was cleared by centrifugation at 30 000 g for 90 min. The supernatant was kept and supplemented with PEG 20 000 until a concentration of 2.8% w/v was reached and subsequently centrifuged at 20 000 g for 5 min. The supernatant was recovered again and further PEG 20 000 added to adjust the percentage to 4.2% w/v before centrifugation at 20 000 g for 10 min. The resulting pellet was resuspended in 35 ml of buffer A and carefully layered on a 25 ml sucrose cushion (10 mM HEPES-KOH pH 7.5, 500 mM KCl, 25 mM Mg(CH_3_COO)_2_, 1.1 M sucrose, 0.5 mM EDTA, 1 mM DTT) before centrifugation at 40 000 rpm for 16 h at 4°C in a T70i rotor (Beckman Coulter). The ribosome pellet was washed twice with 5 mL of buffer E (10 mM HEPES-KOH pH 7.5, 100 mM KCl, 10 mM Mg(CH_3_COO)_2_, 0.5 mM EDTA, 1 mM DTT) before resuspension in 1.5 ml of buffer E. The resuspensions were evaluated by Nanodrop spectrophotometry and approximately 200 AU layered on top of a 7–30% sucrose gradient. Gradients were centrifugated at 17,100 rpm for 16 h 40 min at 4°C in an SW32 rotor (Beckman Coulter). Sucrose gradient tubes were fractionated with a Piston fractionator (Biocomp). Samples corresponding to 70S ribosomes were collected, pooled and the magnesium concentration was adjusted to 25 mM. Subsequently, a final PEG precipitation was performed by addition of PEG 20 000 up to 4.5% w/v and centrifugation at 20 000 g for 12 min. Ribosomal pellets were resuspended in buffer G (10 mM HEPES-KOH pH 7.5, 50 mM KCl, 10 mM NH_4_Cl, 10 mM Mg(CH_3_COO)_2_, 1 mM DTT) and flash frozen in liquid nitrogen until use for biochemical assays or structure determination by cryo-EM.

### Complex assembly and cryo-EM grid preparation

*Staphylococcus aureus* 70S ribosomes were placed on ice, diluted to decrease magnesium concentration to 1 mM, and incubated for 10 minutes at 37°C. *S. aureus in vitro* transcribed *spa* mRNA was unfolded by heating at 80°C for 90 seconds, followed by incubation on ice for 1 min and subsequently at room temperature for 5 min. A mixture of 300 nM *S. aureus* 70S ribosomes, *spa* mRNA, and non-aminoacylated initiator tRNA^fMet^ were mixed at a 1:5:5 molar ratio and incubated for 10 minutes at 37°C. Subsequently, non-aminoacylated tRNA^Lys^ was added in the same molar ratio as the initiator tRNA, and the magnesium concentration was adjusted to 15 mM before a final incubation step of 10 minutes at 37°C.

Three microliters of the assembled *S. aureus* 70S initiation complex was applied to carbon-coated glow-discharged Quantifoil R 2/2 300-mesh copper grids. Grids were blotted and plunge-frozen using a Vitrobot Mark IV (Thermo Fisher Scientific) at 95% humidity and 10°C.

Cryo-EM data were collected on the in-house Glacios transmission electron microscope (Thermo Fisher Scientific) at the CBI operating at 200 kV and equipped with a K2 direct electron detector camera. Images were acquired at a nominal magnification of 45 000 x, corresponding to a calibrated pixel size of 0.9 Å.

### Image processing and model building

All image processing was performed in CryoSPARC v4.7. Preprocessing steps included motion correction, contrast transfer function (CTF) estimation, and particle picking, resulting in 177,831 particles, which were extracted. After 2D classification, 157,007 particles were selected for ab initio 3D reconstruction and subsequent 3D classification. Five distinct classes were obtained, corresponding to: (1) non-rotated 70S with E-site tRNA, (2) rotated 70S with A-site tRNA, (3) rotated 70S with A- and P-site tRNAs, (4) 50S subunit, and (5) junk. Classes 2 and 3 were merged for refinement due to the presence of clear density for the Shine-Dalgarno–anti-Shine-Dalgarno helix and refined by Homogeneous refinement. To identify overrepresented views in the reconstruction, the software UVE (Urzhumtseva, L., 2024) was used. These views were then removed using an updated version of the software (to be published). A final particle stack of 21,621 particles were refined by non-uniform refinement reaching 2.9 Å resolution, as estimated by Fourier Shell Correlation (FSC). For visualization, sharpening was applied using EMReady with default parameters. A schematic overview of Image processing is shown in supplemental Figure S6. The statistics are reported in Table S3.

For model building, the structure of the *S. aureus* 70S initiation complex (PDB: 6YEF (88)) was used as starting atomic model and fitted into the map using the “Fit in Map” tool in UCSF Chimera software. The atomic model was manually inspected and adjusted in Coot v0.9.8.95. The mRNA sequence 5’-GGGGGUAUUAAUUUGAAA-3’ and the anti-SD sequence 5’-GGAUCACCUCCUUU-3’ from the 16S rRNA were built and fitted using RNA tools in the Coot software. The tRNA^Lys^ at the A-site was modeled by rigid-body placement of a tRNA^fMet^, followed by base modifications using the “Mutate” tool to convert it to tRNA^Lys^. The model was refined using real-space refinement in the Phenix v1.21 software, with iterative cycles of manual adjustment in the Coot software.

## Acknowledgments

We are grateful to Dr. Jose Refugio Jaramillo (IBMC, Strasbourg) for his technical assistance and continuous support throughout various aspects of the project. We thank Prof. Alexander Mankin (University of Illinois, Chicago) for his ongoing insights into the methodology, his close engagement with the progression of our results, and his valuable advices. We also thank Prof. Wolfgang R. Hess (University of Freiburg) for helpful discussions on the overall project. We acknowledge Isabelle Caldelari and David Lalaouna for their fruitful discussions. We thank Sasha Ballet, Tan Dat Truong, Léo Fréchin & Charles Barchet for IT support, and Alexandre Durand & Niels Maréchal for cryo-EM support at the integrated structural biology platform of the CBI. This work was supported by the French National Research Agency ANR (SaRNAmod: ANR-21-CE12-0030-01, SatRNAsPG ANR-24-CE11-7652 to SM, IntRNAReg ANR-23-CE12-0041-01 to PR). This work of the Interdisciplinary Thematic Institute IMCBio, as part of the ITI 2021-2028 program of the University of Strasbourg, CNRS and Inserm, was supported by IdEx Unistra (ANR-10-IDEX-0002), and by SFRI-STRAT’US project (ANR 20-SFRI-0012) and EUR IMCBio (ANR-17-EURE-0023) under the framework of the French Investments for the Future Program. The electron microscope facility at the CBI/IGBMC was supported by the Region Grand Est, FEDER, the French Infrastructure for Integrated Structural Biology (FRISBI) ANR-10-INBS-0005 / France 2030 program, EquipEx+ France-Cryo-EM (ANR-21-ESRE-0046) and Instruct-ERIC.

## Author contributions

Project conceptualization, SM, MPK and PR; methodology, MPK, SM, RBC and BPK; investigation, MPK, RBC, MK and ME; analysis, MPK and BCWM; cryo-EM analysis, RBC; visualization, MPK, RBC and SM; funding acquisition, SM, PR and BPK; supervision, SM, PR and BPK; writing – original draft, MPK, SM and PR; writing – review & editing, MPK, SM, PR and BPK, with inputs from everyone.

## Data availability

RNA sequencing data (Ribo-seq, Ribo-Ret and total RNA-seq) applied in this paper is available in the Gene Expression Omnibus (GEO) database under accession numbers GSE299221 and GSE299222.

## Supplemental information

Scheme S1. sORFs discovery pipeline, related to Methods.

Table S1. List of newly discovered sORFs, related to Figure 1.

Table S2. List of oligonucleotides used in this study, related to Methods.

Table S3. Cryo-EM data processing statistics, related to Methods.

---

**Supplemental Scheme S1.**
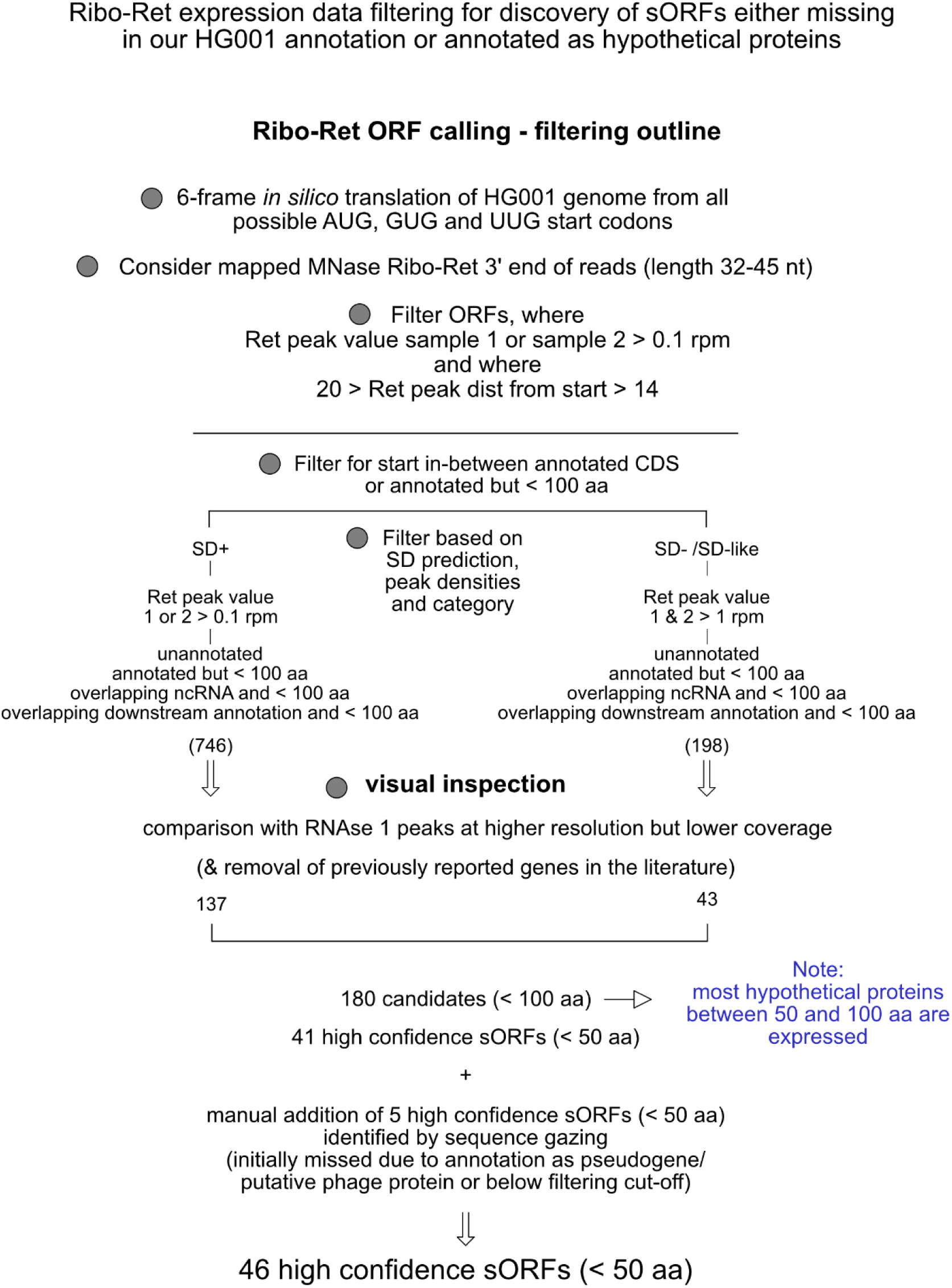
Ribo-Ret expression data filtering scheme for sORF discovery in *S. aureus* HG001, related to Methods and Figure 1. Schematic flow chart representation of the analysis performed to identify sORFs by filtering of mapped Ribo-Ret expression data. Initially, all possible sORFs from AUG, GUG or UUG start codons were *in silico* translated into candidate sequences. Subsequently, they were either kept or discarded based on whether Ribo-Ret density peaks were adequately positioned in respect to the putative start sites and above indicated rpm expression thresholds. Key decisions during the analysis with this filtering approach are indicated by grey circles. Differential criteria considered for filtering, such as predicted SD-motif presence are designated on top of each branch of the decision tree. The number of candidate-sORFs remaining after computational filtering and after visual inspection are indicated accordingly.

**Figure S1.**
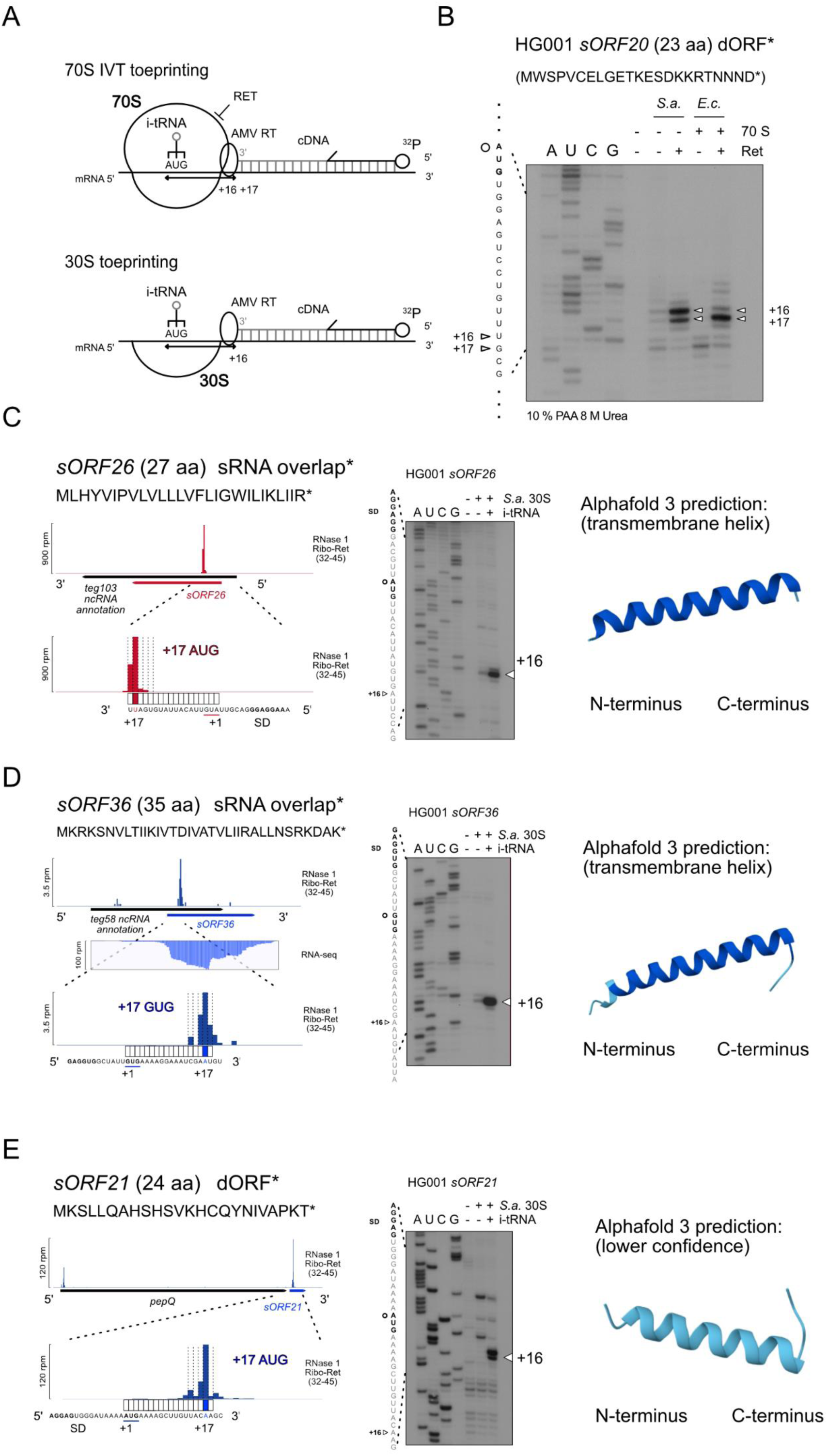
Biochemical validation of initiation complex formation on novel sORF candidate sequences, related to Figure 1. (A) Graphical illustration of toeprinting analysis for monitoring of translation initiation complex formation. Initiation complexes were formed on candidate sORF sequences either with purified *S. aureus* 70S ribosomes, trapped by addition of Ret during *in vitro* translation, or with 30S subunits. Primer extension inhibition analysis of radiolabeled oligos was used to assess the position and efficiency of initiation complex formation. (B) 70S initiation complex toeprinting analysis of candidate *sORF20*, referenced in main figure 1E. Toeprint bands at positions +16 and +17 in respect to the predicted AUG start codon of *sORF20* as well as sequencing lanes and the corresponding nucleotide sequence are indicated accordingly. Toeprints obtained by use of either *S. aureus* or *E. coli* purified 70S are shown. (C, D, E) Ribo-Ret profiles of *S. aureus sORF26* (C)*, sORF36* (D) and *sORF21* (E), displaying ribosomal density peaks and a zoomed focus on their respective initiation sites. Toeprinting analyses indicating 30S initiation complex formation corresponding to each of the Ribo-Ret footprints are shown on the right of each panel, next to a structural model of the designated small proteins as predicted by AlphaFold. The respective small protein sequence is indicated on top of each Ribo-Ret density profile. For each 30S toeprint, sequencing lanes and the corresponding nucleotide sequence are displayed.

**Figure S2.**
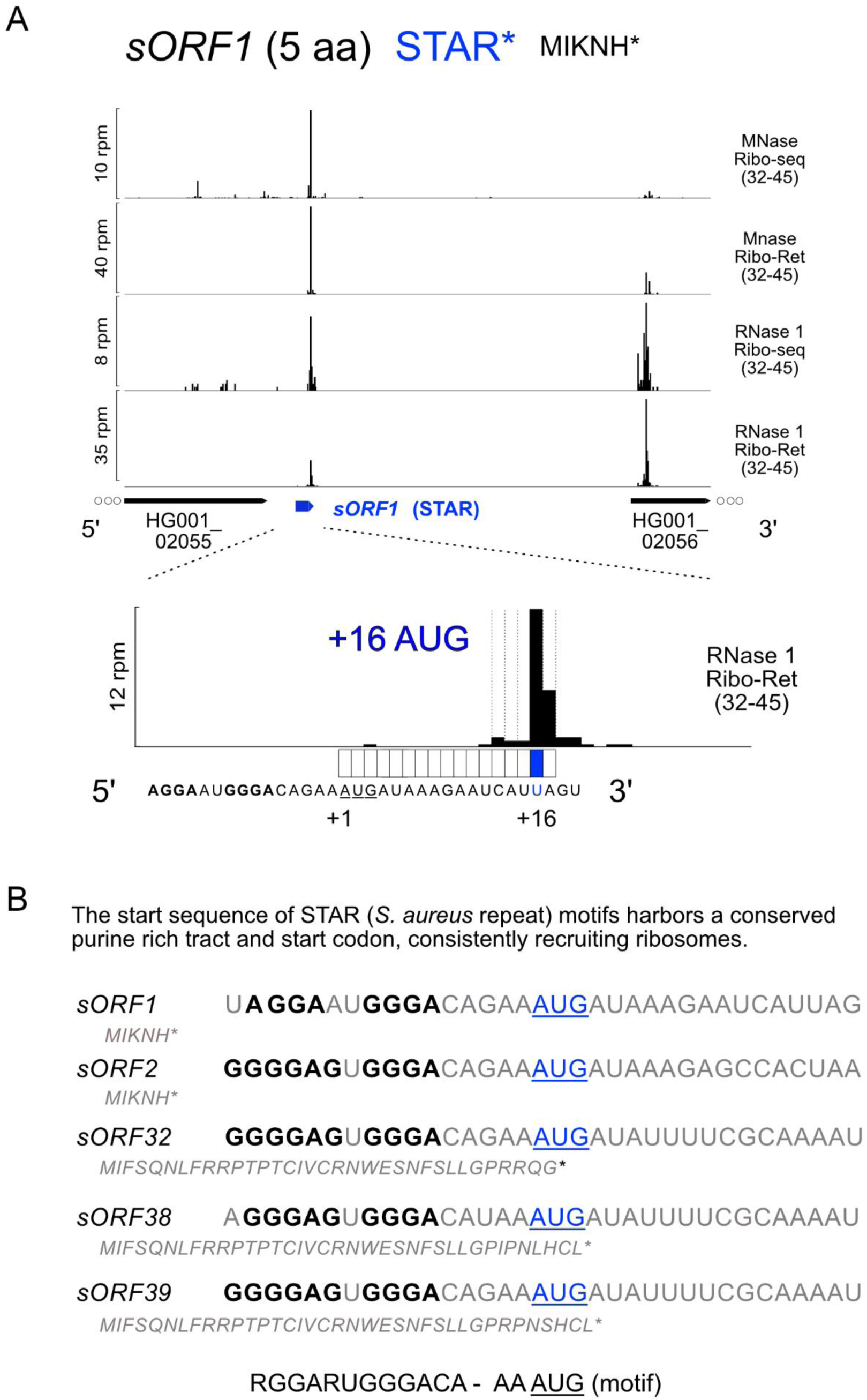
Staphylococcal repeat (STAR) motifs recruit ribosomes for translation initiation at their conserved 5’ sequences, designating sORFs of variable length, related to Figure 1. (A) Exemplary Ribo-Ret profiles of *sORF1*, corresponding to a 5 aa short sORF, whose initiation site is one of several conserved STAR motif 5’ sequences. The exact position of the Ribo-Ret footprint is shown on a zoomed view at the bottom, highlighting 3’ end density at position +16 in respect to an AUG start codon. (B) Sequence comparison of the RBS motifs of STAR sequences, displaying Ribo-Ret densities and designating variable sORFs, expressed in our experimental conditions. The selected start codons (blue) with appropriately spaced purine-rich tracts (bold) are highlighted in the alignments and a corresponding sequence motif is displayed below. The different small proteins designated by the five STAR sequences shown, are indicated below their respective sORF identifiers, in grey.

**Figure S3:**
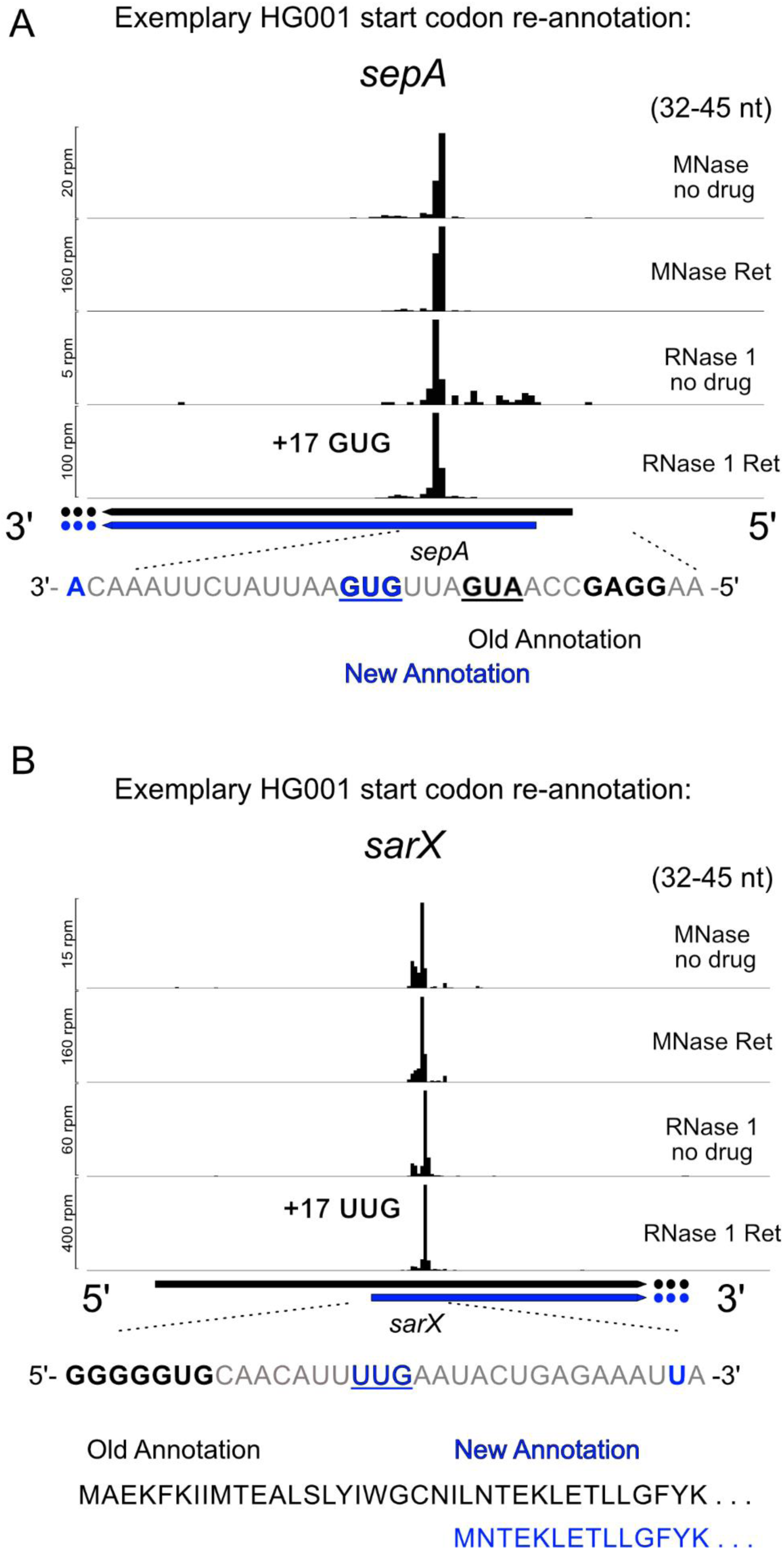
High-resolution Ribo-Ret facilitated improved genome annotation, resulting in minor and major start codon re-assignments, related to Figures 1 and 2. (A) Exemplary Ribo-Ret profiles of *sepA*, with density peaks, whose positions designate alternative in-frame start codon selection. An appropriately spaced GUG start codon (blue), was re-annotated compared to the previously annotated AUG start codon (black). This example is representative of several minor re-annotations as a result of improved resolution by RNase 1 Ribo-Ret. (B) Exemplary Ribo-Ret profiles of *sarX*, with density peaks far from the previously annotated putative initiation site, therefore designating a major re-annotation. The previous 22 aa longer N-terminal annotation is indicated in black, while the new annotation from an appropriately spaced UUG start codon with strong Ribo-Ret peak, is shown in blue.

**Figure S4:**
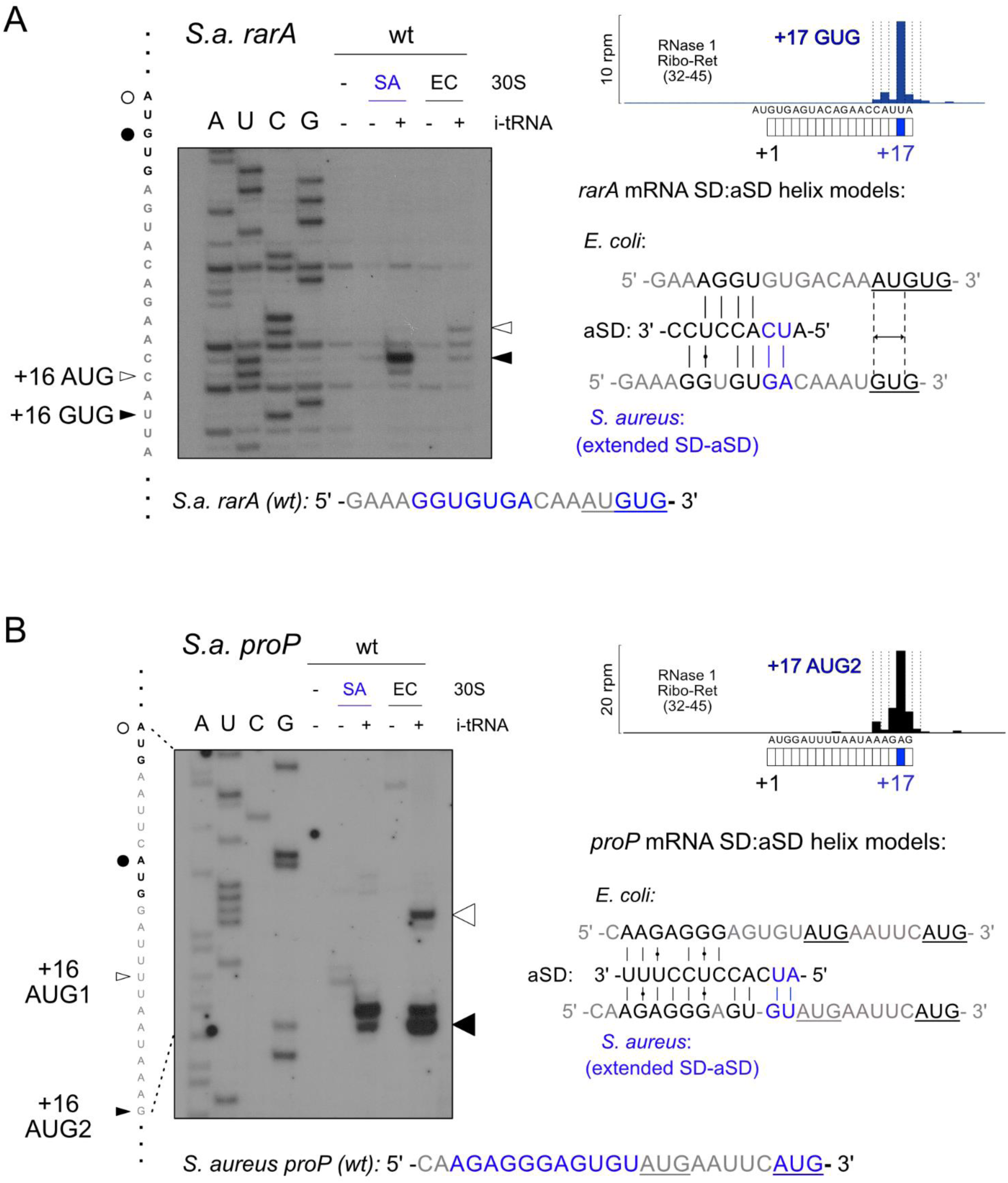
Alternative SD-aSD alignment during translation initiation through extended SD-aSD formation can direct differential start codon selection between *S. aureus* and *E. coli*, related to Figure 3. (A, B) Comparative 30S toeprinting analysis of differential start codon selection by *S. aureus* and *E. coli* ribosomes on the natural *S. aureus* mRNA sequences *rarA* (A) and *proP* (B). For both mRNAs’ SD-motifs, extended SD-aSD interactions are predicted to form. The predicted model for *rarA* is displayed on the right of the toeprint panel, below its RNase 1 Ribo-Ret density profile (A). An extended interaction may differently align the SD and aSD sequence on this particular mRNA. In panel B on the other hand, the model for the SD-aSD interaction on *proP*, shown below its Ribo-Ret density profile, indicates similar alignments between *S. aureus* and *E. coli*. However, the possible extended interactions may preclude initiation at the first of two potential AUG start codons. For both panels, toeprint bands at positions +16 (arrows) in respect to either potential start codon, sequencing lanes and the corresponding nucleotide sequence are indicated accordingly. For both *rarA* as well as *proP*, the full RBS sequence is displayed below the toeprint gel, highlighting the extended SD motif and the in-frame start codon in *S. aureus* in blue.

**Figure S5:**
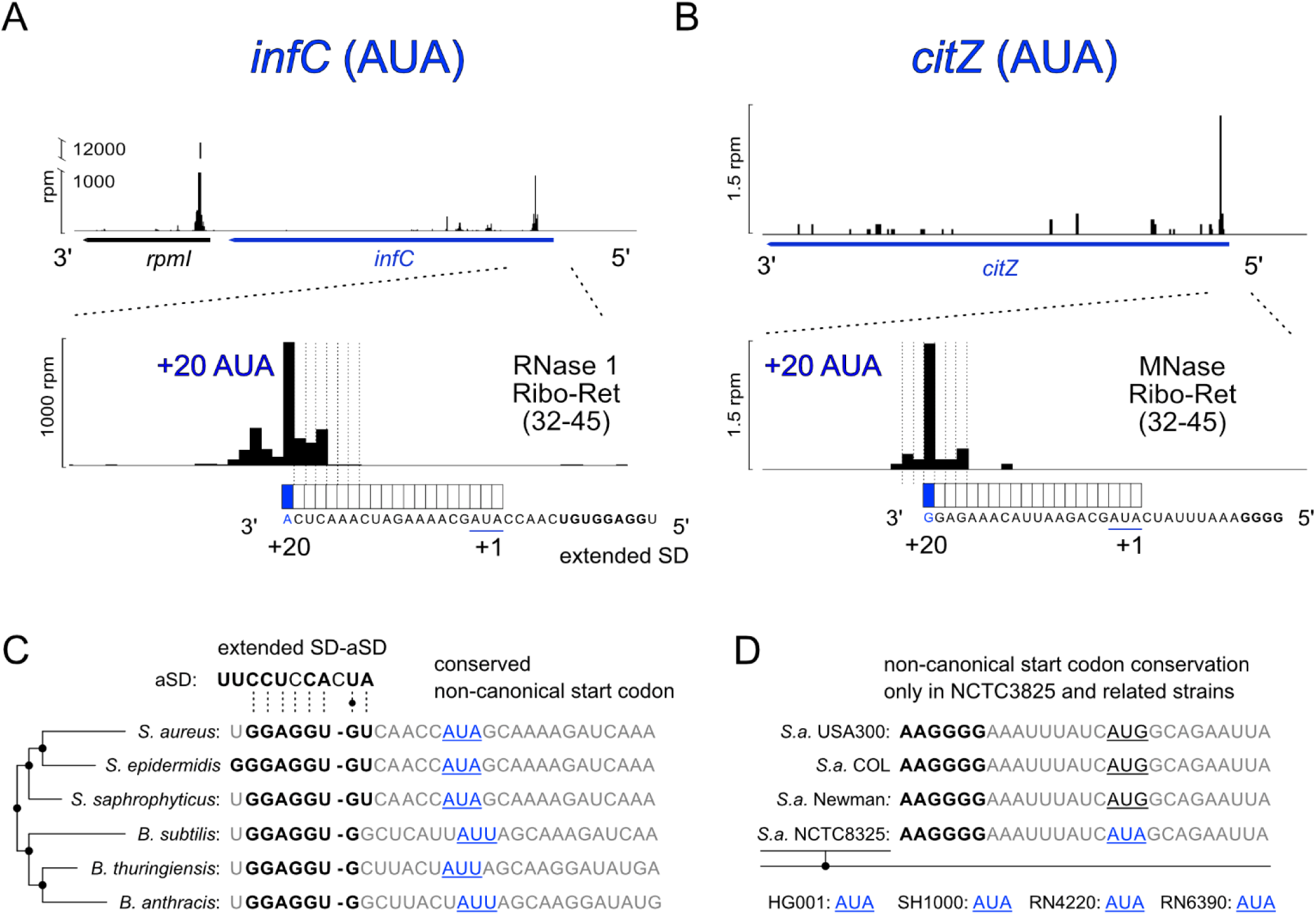
*infC* and *citZ* utilize non-canonical AUA start codons in *S. aureus* HG001 strain, related to Figure 4. (A, B) Non-canonical start codon usage by the *infC* (A) and *citZ* (B) mRNAs is shown, evidenced by their respective Ribo-Ret expression profiles. The Ribo-Ret data type used for visualization is indicated accordingly, e.g., indicating the use of MNase Ribo-Ret data of read lengths 32-45 nt for *citZ* (B) due to low expression values and sequencing coverage of the locus. (C, D) Evolutionary RBS sequence conservation analysis of *infC* (C) and *citZ* (D), as performed by comparison of curated genome information extracted from the SEED database. Potential SD-aSD motif interactions are shown in bold, including putative extended interactions in the case of *infC* (C), while conserved non-canonical start codons are highlighted in blue.

**Figure S6:**
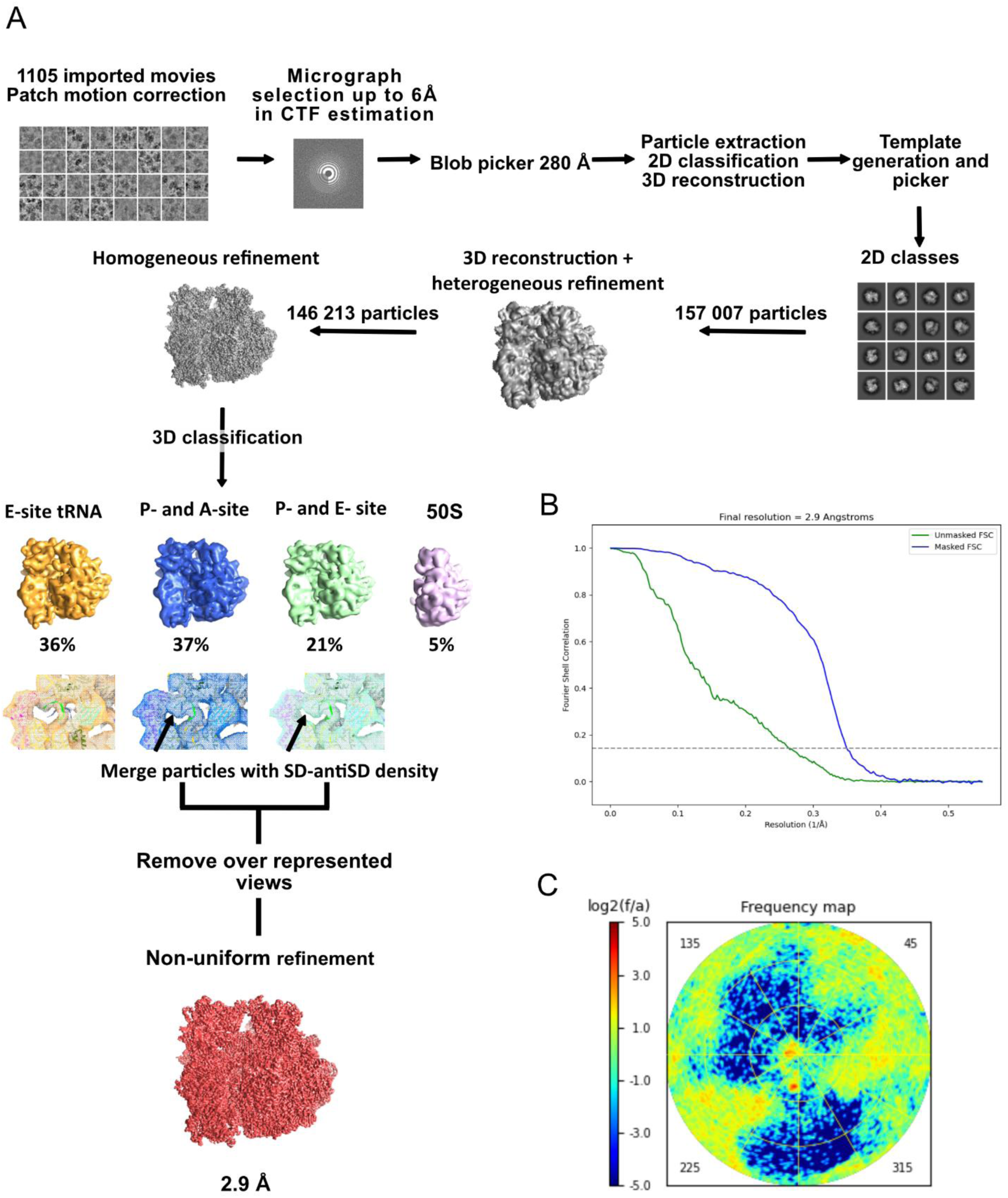
Single particle cryo-EM data processing on *S. aureus* 70S initiation complex, related to Methods and Figure 3. (A). Particle sorting scheme. Arrows in 3D classification indicate presence of signal corresponding to SD-antiSD helix. (B) Estimated resolution based on Fourier shell correlation at 0.143 (dotted line). (C) View distribution from map represented by VUE (89)

**Table S1.**
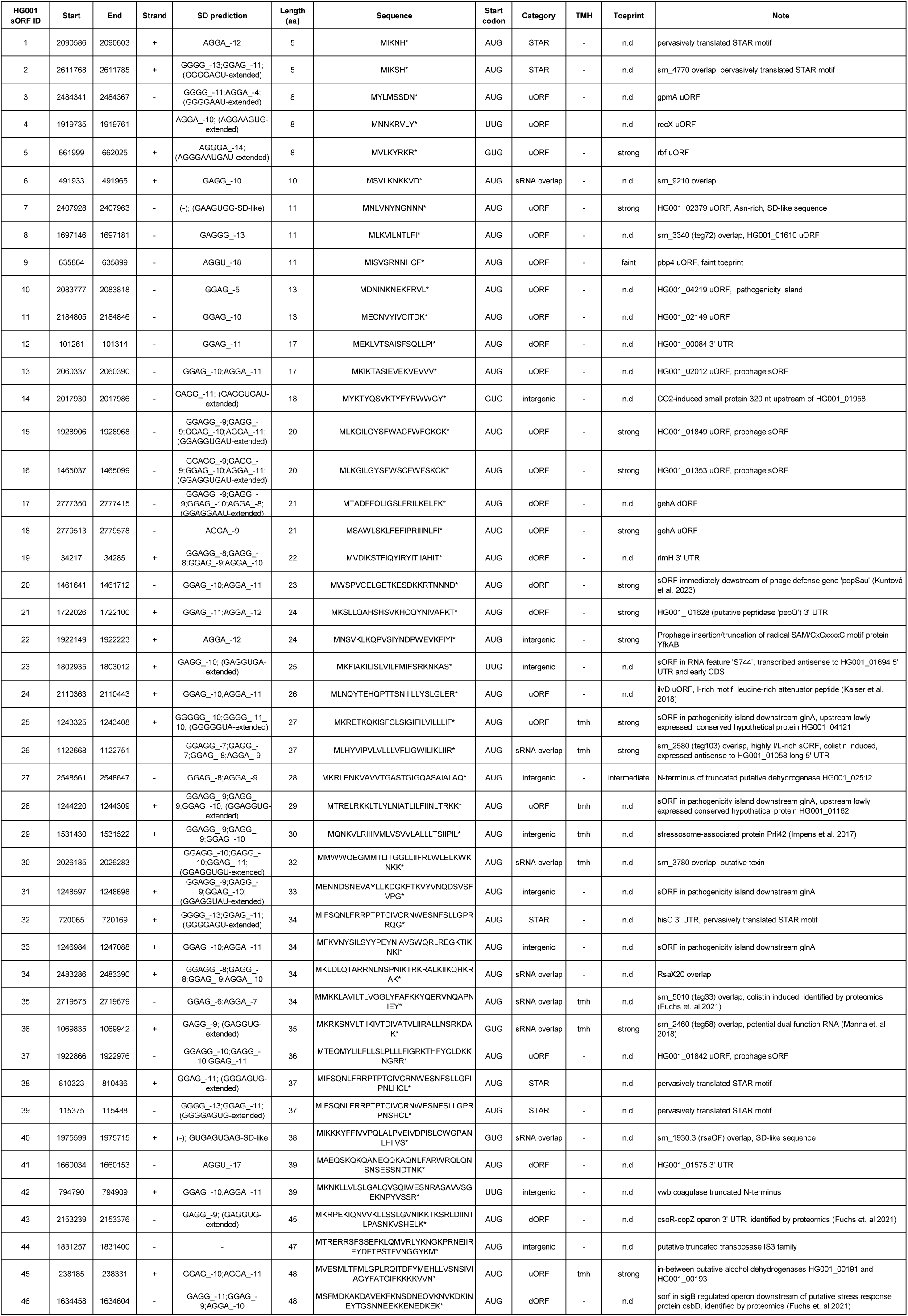
Novel unannotated sORFs identified in *S.a*. HG001.

**Table S2.**
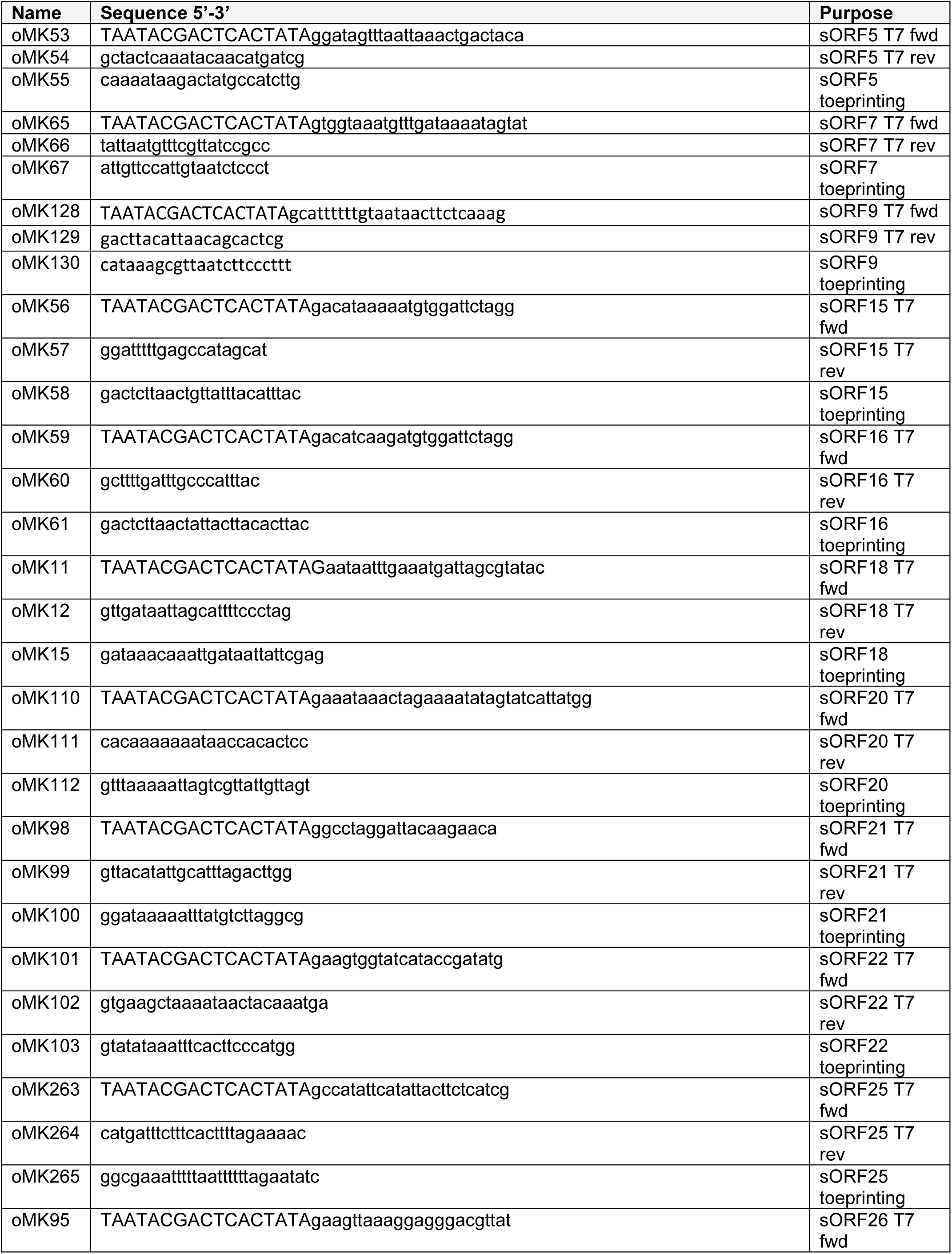
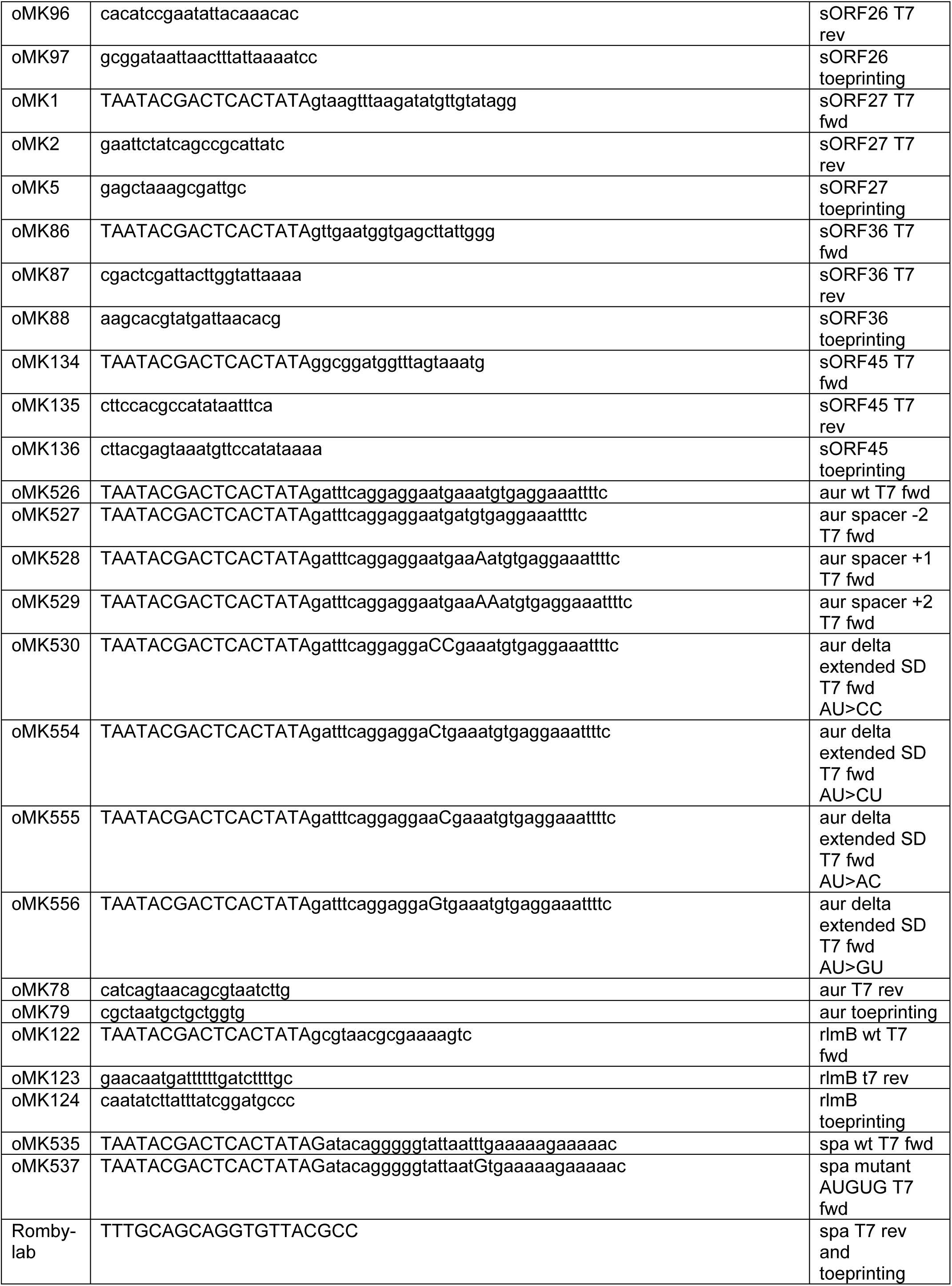
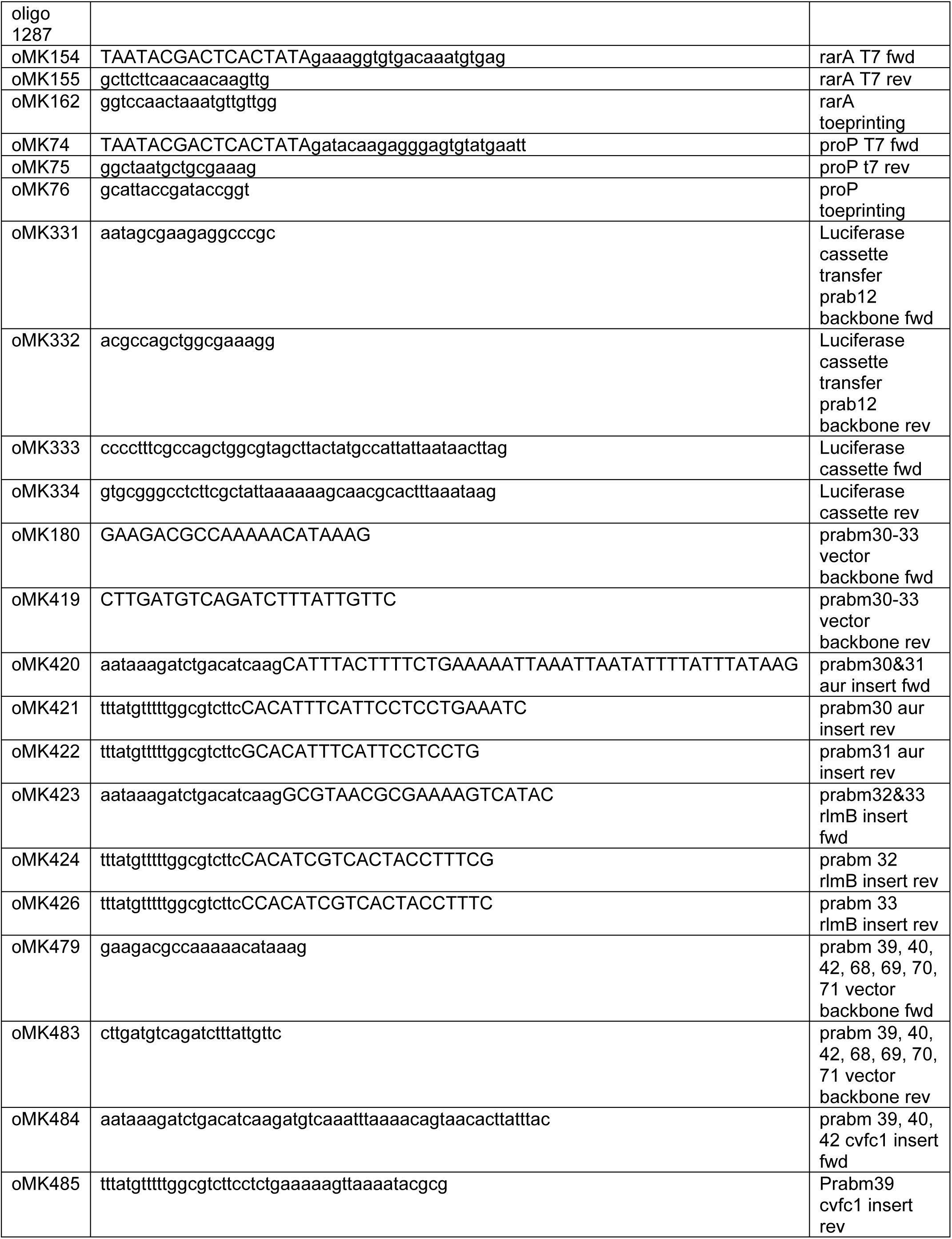
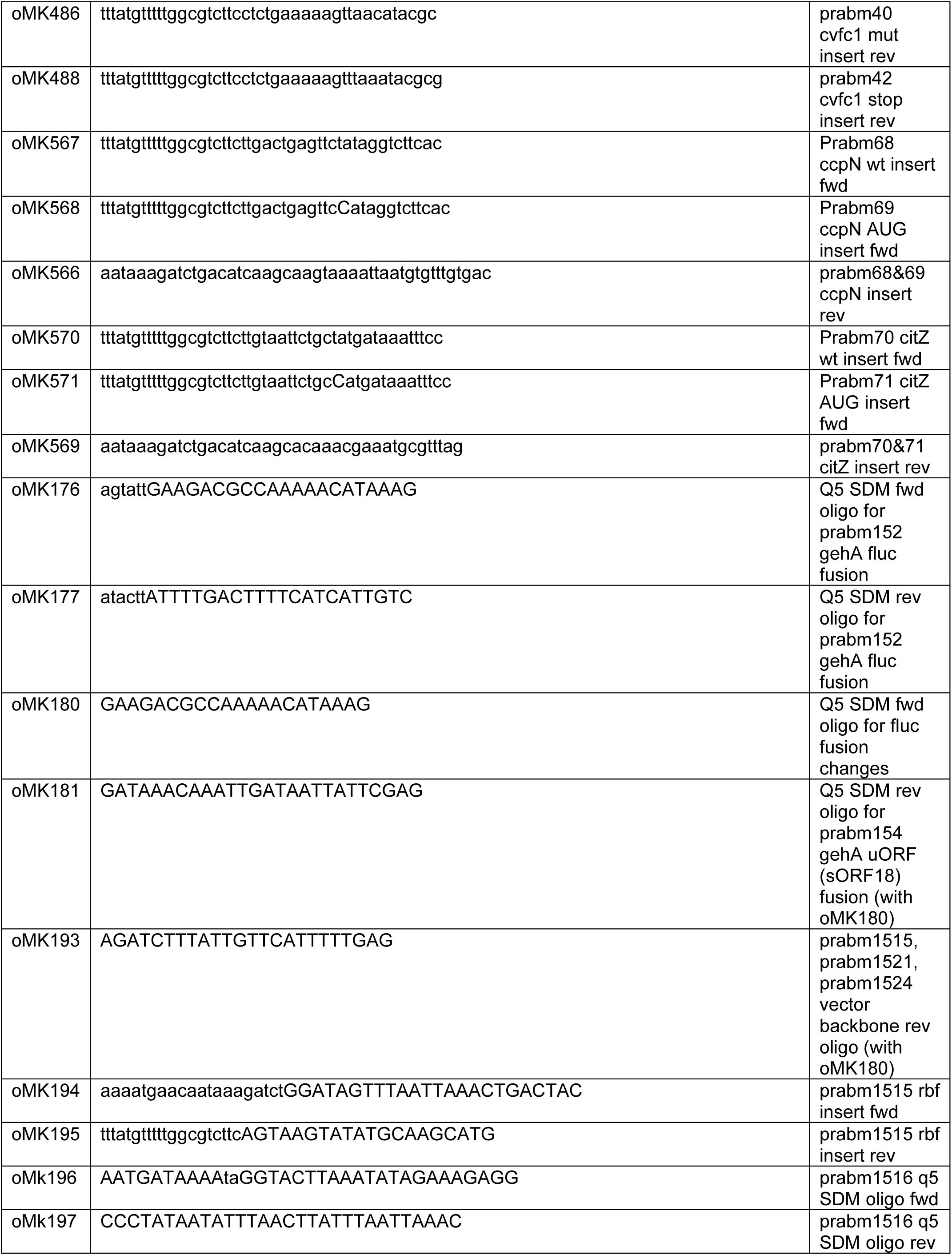
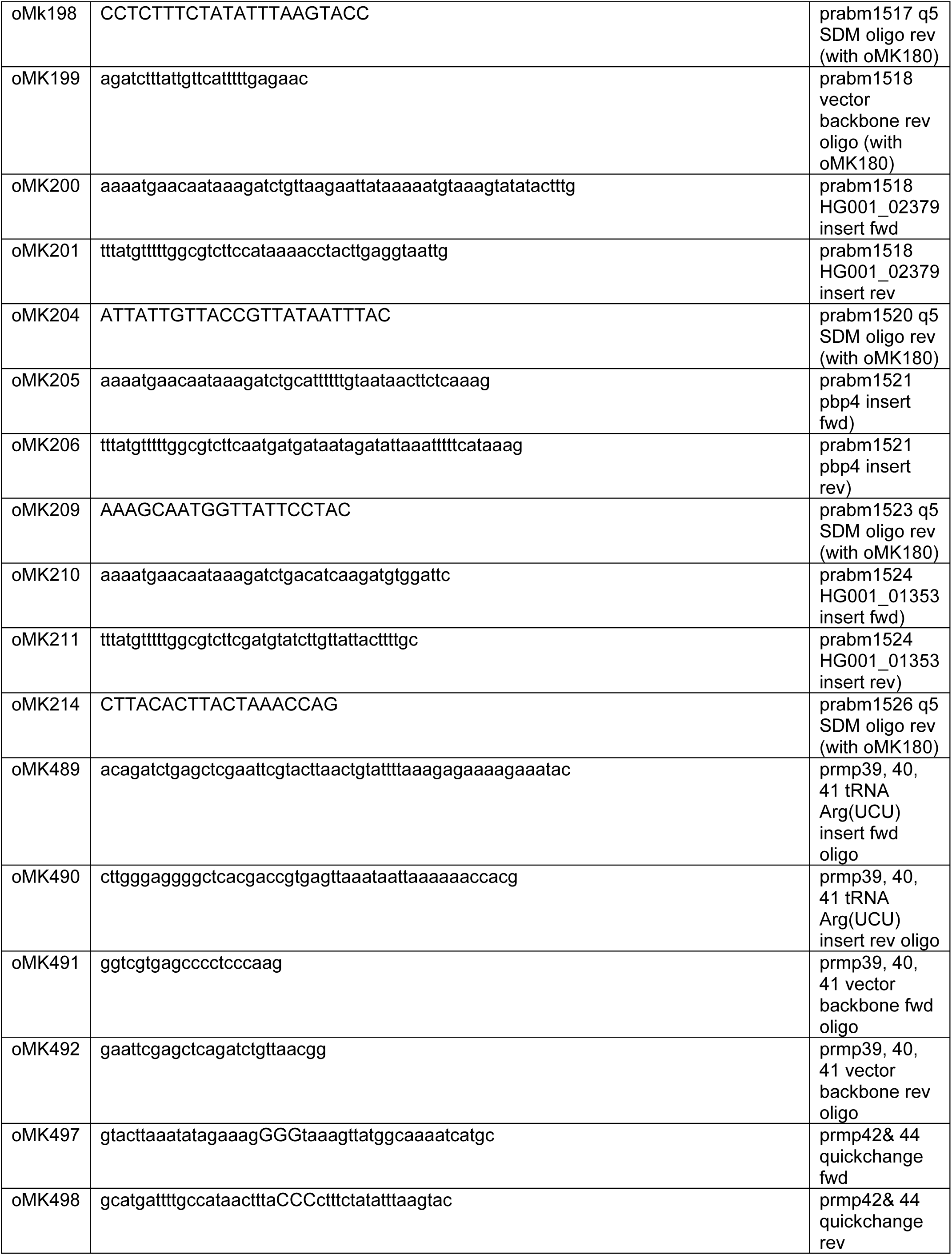
List of oligonucleotides used in this study.

**Table S3.** Cryo-EM data processing statistics, related to Methods.

|  |  |
| --- | --- |
| **Data collection processing** |  |
| Magnification | 45, 000 |
| Acceleration Voltage (KV) | 200 |
| Electron exposure (e$\text{\AA}^{-2}$) | 30 |
| Defocus range ($\mu\text{m}$) | -0.5 to 2.5 |
| Pixel size | 0.91 |
| Simmetry imposed | C1 |
| Initial particle images (no.) | 177 831 |
| Final particle images (no.) | 21 621 |
| Map resolution ($\text{\AA}$) | 2.9 |
| Fourier Shell correlation threshold | 0.143 |
| **Refinement** |  |
| Initial model used (PDB code) | 6YEF |
| Model resolution | 2.9 |
| Fourier shell correlation threshold | 0.5 |
| Map sharpening B factor ($\text{\AA}$) | 0 |
| **Model composition** |  |
| Non-hydrogens atoms | 140814 |
| Protein residues | 5275 |
| Nucleotides | 4634 |
| **B factors ($\text{\AA}$)** |  |
| Protein | 36.6 |
| RNA | 36.0 |
| **Root mean square deviations** |  |
| Bond lengths ($\text{\AA}$) | 0.005 |
| Bonde angles ($\text{\AA}$) | 0.595 |
| **Validation** |  |
| Molprobity score | 1.82 |
| Clashscore | 10.20 |
| Rotamer outliers | 0.07 |
| **Ramachandran plot** |  |
| Favored (%) | 95.79 |
| Allowed (%) | 4.05 |
| Disallowed (%) | 0.15 |

